# Large Freshwater Phages with the Potential to Augment Aerobic Methane Oxidation

**DOI:** 10.1101/2020.02.13.942896

**Authors:** Lin-Xing Chen, Raphaël Méheust, Alexander Crits-Christoph, Katherine D. McMahon, Tara Colenbrander Nelson, Lesley A. Warren, Jillian F. Banfield

## Abstract

There is growing evidence that phages with unusually large genomes are common across various natural and human microbiomes, but little is known about their genetic inventories or potential ecosystem impacts. Here, we reconstructed large phage genomes from freshwater lakes known to contain bacteria that oxidize methane. Twenty-two manually curated genomes (18 are complete) ranging from 159 to 527 kbp in length were found to encode the *pmoC* gene, an enzymatically critical subunit of the particulate methane monooxygenase, the predominant methane oxidation catalyst in nature. The phage-associated PmoC show high similarity (> 90%) and affiliate phylogenetically with those of coexisting bacterial methanotrophs, and their abundance patterns correlate with the abundances of these bacteria, supporting host-phage relationships. We suggest that phage PmoC has similar functions to additional copies of PmoC encoded in bacterial genomes, thus contribute to growth on methane. Transcriptomics data from one system showed that the phage-associated *pmoC* genes are actively expressed *in situ*. Augmentation of bacterial methane oxidation by pmoC-phages during infection could modulate the efflux of this powerful greenhouse gas into the environment.

## Introduction

Bacteriophages (or phages), viruses that infect and replicate within Bacteria, are important in both natural and human microbiomes because they prey upon bacterial hosts, mediate horizontal gene transfer (HGT), alter host metabolisms, and redistribute bacterially-derived compounds via host cell lysis ^1^. A phenomenon that has recently come to light via cultivation independent studies is the prominence of phages with genomes that are much larger than the average size of ∼55 kbp predicted based on current genome databases ^2^. The newly reported genomes range up to 735 kbp in length and are reported to encode a diversity of genes involved in transcription and translation as well as genes that may augment host metabolism ^2^. Augmentation of bacterial energy generation by auxiliary metabolic genes (AMGs) has been reported for phages with smaller genomes. For example, some encode photosynthesis-related enzymes ^3^, some deep-sea phages have sulfur oxidation genes ^4^, and others that infect marine ammonia-oxidizing Thaumarchaeota harbor a homolog of ammonia monooxygenase subunit C (i.e., *amoC*) ^5,6^. Unreported to date is the role of phages involved in the oxidation of methane, a greenhouse gas that is 20-23 times more effective than CO_2_ ^7^. Biological transformation of methane is largely driven by microorganisms, including aerobic methanotrophs belonging to Alphaproteobacteria and Gammaproteobacteria and Verrucomicrobia that use soluble methane monooxygenases (sMMO) and/or particulate methane monooxygenases (pMMO) ^8^. The pMMO, the predominant methane oxidation catalyst in nature, is a 300 kDa trimeric metalloenzyme ^9^ that converts methane to methanol in the periplasm ^8,10^. The pMMO is encoded by the *pmoCAB* operon ^11^ and some bacterial genomes encode multiple *pmoCAB* operons as well as additional copies of *pmoC* that appear to be essential for growth on methane ^12^.

We considered the possibility that phages infecting methanotrophs could directly impact methane oxidation and thus methane release. Phages with very large genomes were recently reported from a man-made lake that covers a deposit of methane-generating tailings from an oil sands mine in Canada ^2^. Here, we searched the unreported phage genome fragments from this lake for genes involved in methane oxidation. We identified four sets of assembled fragments comprising draft genomes that encoded the enzymatically critical *pmoC* subunit. Hereafter, we refer to these phages as pmoC-phages. We also investigated the metagenomic datasets from Crystal Bog, Lake Mendota, and Trout Bog Lake, freshwater lakes in Wisconsin, USA, that are known sources of sediment-derived methane ^13–15^ and found examples of *pmoC* on a subset of phage genome fragments from all three ecosystems. Of the 16 pmoC-phage genomes, twelve were manually curated to completion, enabling verification that they do not encode *pmoA* or *pmoB* genes. All complete and partial genomes are > 165 kbp in length. Microbial communities from all three lakes are known to contain proteobacterial methanotrophs ^16^, some of which we infer are the hosts for these phages. We suggest that pmoC-phages may play important roles in the methane cycle.

## Results

### Active methane oxidation in a methane-generating tailings lake

Oil sands (bituminous sands) deposits are mined for petroleum and generate large volumes of waste materials that produce methane, hydrogen sulfide, and ammonia. The oil sands pit lake of Base Mine Lake (BML) in Alberta (Canada) was constructed by placing a layer of water over a tailings deposit, with the long-term goal of developing a lake ecosystem supported by a stable oxic zone. The oxic zone would permit the oxidation of methane, ammonia and hydrogen sulfide. The lake is characterized by high concentrations of methane and ammonia (up to 253 μM and 73.5 μM, respectively; Fig. 1a, Supplementary Table 1), especially in the hypolimnion layer (the lower layer of water in a stratified lake) and at the tailings-water interface ^17,18^. DNA-SIP (stable isotope probing) analyses ^19^ indicate that the methane was produced from fermentation products by archaea in the tailings. We observed a significant sink of methane in the hypolimnion layer (Fig. 1a). For example, in 2015 and 2016, either oxygen or methane (or both, in some samples) were consumed completely, and the indigenous bacterial communities likely used methane as a primary carbon source for growth.

**Fig. 1.**
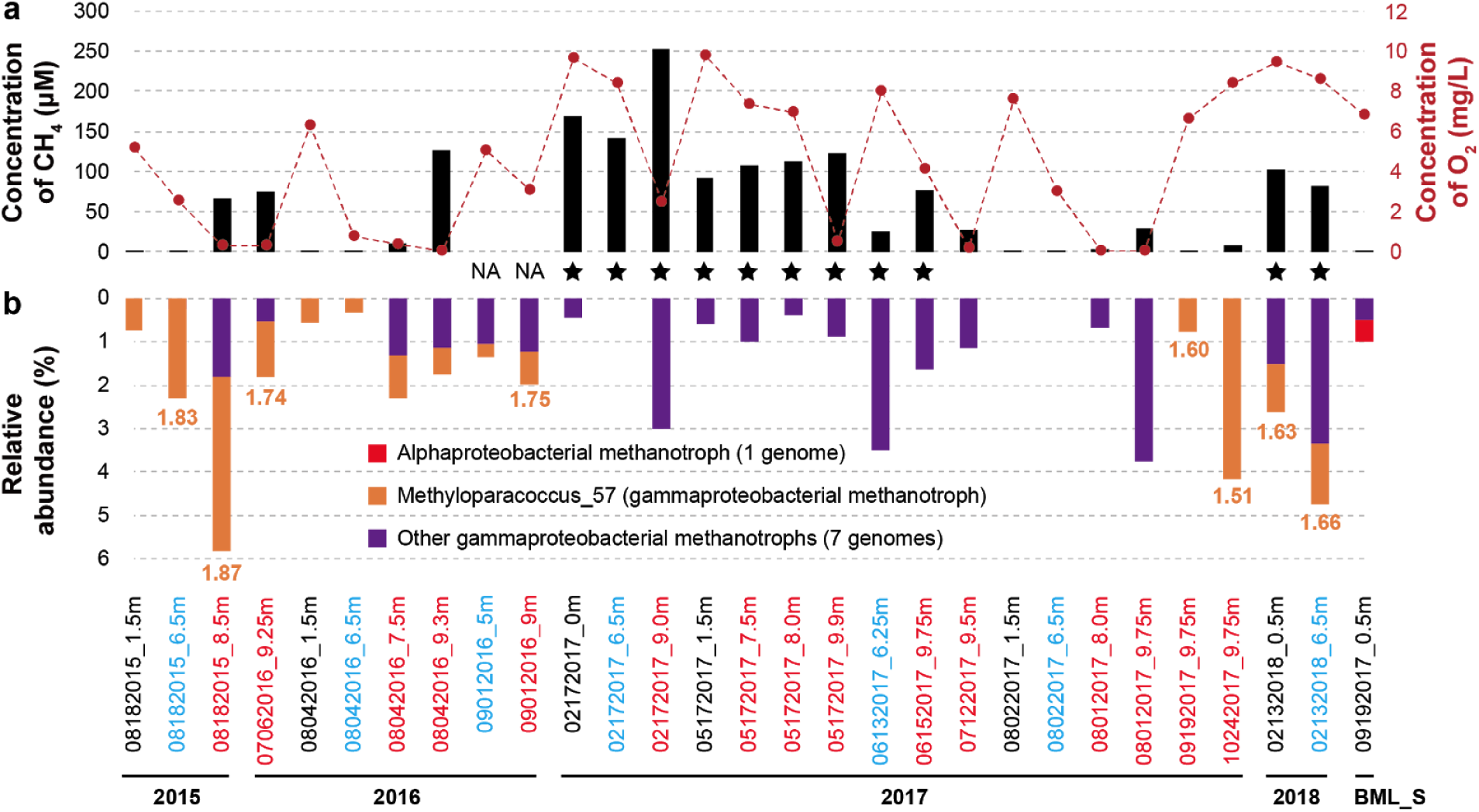
Geochemical and biological evidence for methane oxidation in BML and BML_S samples. (a) The concentration of methane and oxygen determined at different depths (see names; epilimnion in black, metalimnion in blue and hypolimnion in red) at each sampling time point. Samples in which methanotrophs are inferred to be inactive are indicated by stars. NA, not available. (b) The relative abundances of alphaproteobacterial and gammaproteobacterial methanotrophs detected in each sample. The iRep values (orange font) indicative of the growth rates of Methyloparacoccus_57 are shown where the genome coverage was ≥ 5X. The iRep values for other methanotrophs are provided in Supplementary Fig. 2.

Genome-resolved metagenomics was used to identify microorganisms involved in methane oxidation in the 28 BML water samples and one sample from the creek that supplies water to the lake (BML source, BML_S) (Supplementary Fig. 1). We reconstructed genomes of eight gammaproteobacterial methanotrophs that were more abundant in the hypolimnion than the upper layers, and one alphaproteobacterial from BML_S (Fig. 1b, Supplementary Figs. 2 and 3, Supplementary Table 2). Genes encoding sMMO and/or pMMO were detected in these genomes, and some contain more than one copy of *pmoCAB* operon and also standalone *pmoC* (Supplementary Figs. 4 and 5, Supplementary Tables 2 and 3). The most frequently detected methanotroph in BML communities, Methyloparacoccus_57 (Supplementary Fig. 6), shares 96.3% 16S rRNA gene sequence similarity with that of *Methyloparacoccus murrellii* strain R-49797^T 20^. Methyloparacoccus_57 may be a key player in methane oxidation (Fig. 1a) as it had a higher growth rate (iRep values of 1.51-1.87) than any other aerobic methanotrophs (iRep values of 1.32-1.61) coexisting in the communities, especially in the hypolimnion layers of 2015 and 2016 (Fig. 1b and Supplementary Fig. S2). Methane accumulated in lake samples collected from February to June of 2017 and in February of 2018 despite the availability of oxygen (Fig. 1a, Supplementary Table 1), suggesting that low temperatures inhibited the activity of methanotrophs. Reanalyses of published oil sands datasets from Canada detected Methyloparacoccus_57 in other sites ^21–24^ (Supplementary Information), suggesting their potentially significant role in the sink of methane in oil sands related habitats of Canada.

### Phages with standalone *pmoC* genes

Genomes of large phages from the BML samples were previously reported ^2^, but many other phage genome fragments remained to be analyzed. We searched the full set of phage genomes for genes that could contribute to methane oxidation and found *pmoC* genes that shared > 86% amino acid identity to that of published bacterial methanotrophs (Supplementary Table 4). Most of the phage (pmoC-phage) scaffolds ended at the *pmoC* gene, apparently because the assembly was confounded by very similar *pmoC* genes encoded in coexisting bacterial genomes. Manual scaffold extension confirmed no gene encoding *pmoA*/*pmoB* located nearby (see Supplementary Figs. 7 and 8 for example). One of these pmoC-phage genomes (i.e., BML_4) was curated to completion (circularized; see below) to confirm the absence of *pmoA*/*pmoB*.

Reanalysis of the published oil sands datasets (Supplementary Information) detected one pmoC-phage scaffold (TP6_1) in a Suncor tailings pond sampled in 2012 ^23^. Additionally, phages similar to TP6_1 and BML_3 were detected in two other samples from Alberta, i.e., TP_MLSB collected in 2011 ^23^ and PDSYNTPWS collected in 2012 ^22^. From PDSYNTPWS, we curated a phage genome without *pmoC* (referred to as “PDSYNTPWS_1”), which is 99% similar to BML_3 (64% and 75% aligned fraction, respectively). Our reanalysis of published ^13^CH_4_-based DNA-SIP ^22^ data detected PDSYNTPWS_1 in the heavy DNA-SIP fraction. This fraction was dominated by Methyloparacoccus_57, suggesting that this bacterium oxidized ^13^C-enriched methane and could have been the host for the phage.

To test for phage-associated *pmoC* in other lakes reported to emit methane ^13–15^, we searched our previously published metagenomic datasets from Lake Mendota (LM), Crystal Bog (CB) and Trout Bog Lake (TBL) in Wisconsin (USA). HMM-based searches detected *pmoC* on phage scaffolds from all the three lakes (Supplementary Table 5), suggesting the potentially wide distribution of related phages in habitats with methane. The LM, CB and TBL datasets with pmoC-phage scaffolds were reanalyzed in detail (Methods).

We confirmed the high similarity of the bacterial and phage-associated PmoC predicted from all datasets to PmoC of previously described alphaproteobacterial and gammaproteobacterial methanotrophs (Supplementary Table 6). Alignment of these PmoC with references from well-known bacterial methanotrophs ^25^ confirmed the presence of the residues necessary for the copper-binding site, i.e., Asp^156^, His^160^, and His^173^ (Fig. 2a, Supplementary Fig. 9) ^9^ and required for O_2_ binding and methane oxidation ^26^. Interestingly, the bacterial and phage-associated PmoC sequences were generally very similar in the central membrane- and periplasmic associated portions, but divergent at the cytoplasmic N- and C-termini. The *pmoC* genes in three of the pmoC-phages were fragmented and one contained only the C-terminus (Fig. 2a, Supplementary Fig. 10). Interestingly, the *pmoC* from CB_5 exhibited within-population variation as a subset of phages lacks the central region where the active site is located.

**Fig. 2.**
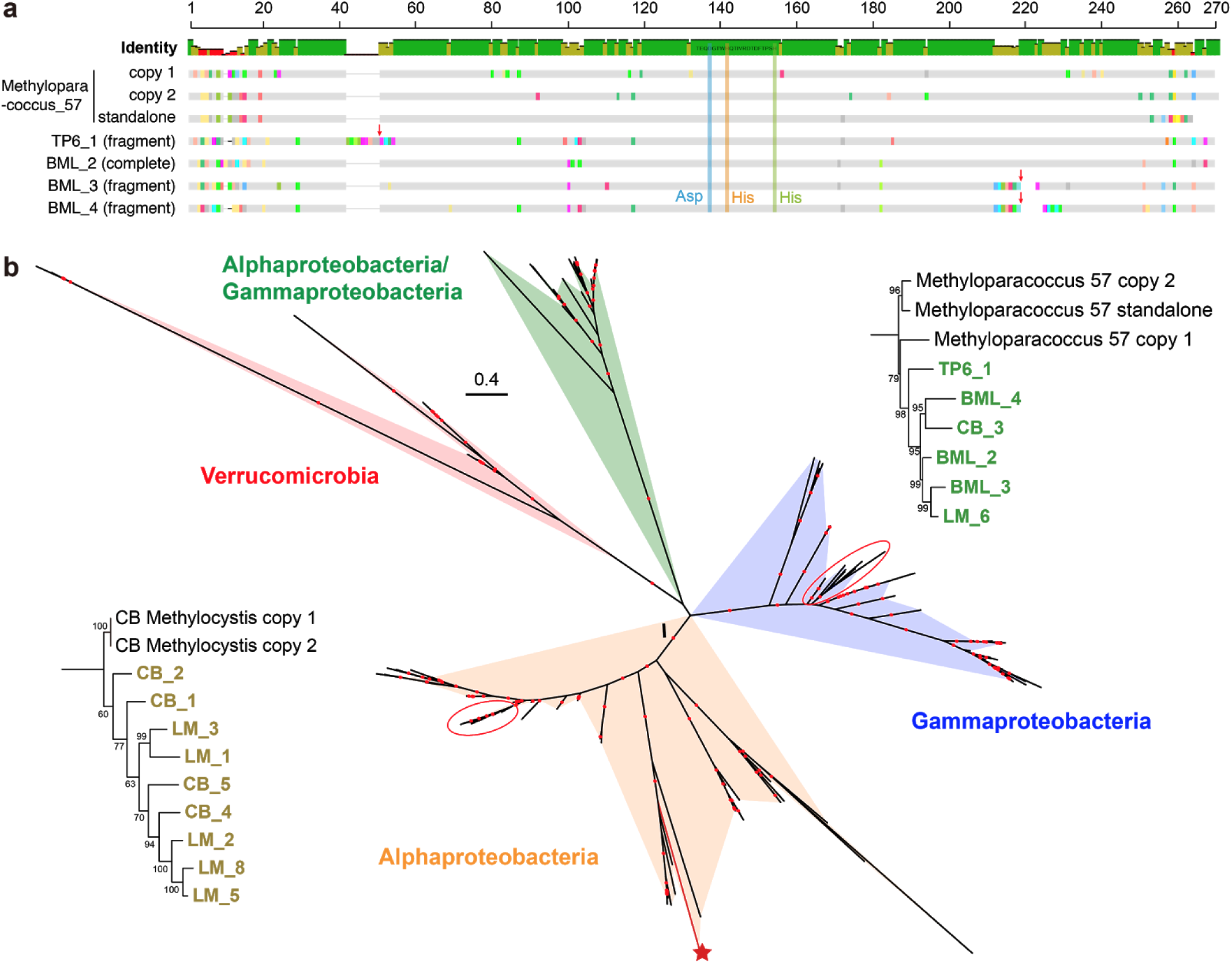
Bacterial and phage-associated PmoC. (a) Alignment of some bacterial and phage-associated PmoC sequences. The three residues in PmoC for copper ion coordination are highlighted (near the middle of the sequence). The *pmoC* genes of phages TP6_1, BML_3 and BML_4 are fragmented; both pieces are shown (red arrows). See Supplementary Fig. 9 for the alignment of all phage-associated and bacterial PmoC sequences. (b) Phylogenetic analysis of bacterial and phage-associated PmoC. Note that no high-quality genome of phage with *pmoC* was reconstructed from TBL. The colored regions show the clades of published and newly reported bacterial sequences. The phylogenies of phage-associated PmoC are shown in detail. The CB *Methylocystis* has a standalone copy of *pmoC* (without *pmoA* or pmoB nearby), the product (red star) of which is distantly related to the two operon-based copies. The PmoC fragments from three BML pmoC-phages were respectively concatenated as one. The bootstraps are indicated by red dots when ≥ 70.

Phylogenetic analyses showed that, regardless of the sampling site, within each taxonomic clade (Alpha- or Beta-) the phage-associated PmoC often clustered together (Fig. 2b, Supplementary Fig. 11). Moreover, the phage-associated PmoC was always more similar to the PmoC of bacterial methanotrophs coexisting in the communities than to the published bacterial PmoC. Interestingly, the total abundance of phage-associated *pmoC* was higher than that of bacterial *pmoC* in some samples (Supplementary Fig. 12).

### Genomic features and taxonomy of pmoC-phages

A total of 22 unique pmoC-phage scaffolds with sequencing coverage > 20X were selected for manual curation to completion (Supplementary Tables 2 and 6) and 18 were completed (no gaps and circular; Table 1). In addition, one partial and four complete genomes of closely related phages but without *pmoC* were manually reconstructed for comparison (see below). The phage genomes are 159-527 kbp in length (GC content: 32-44%), encode between 224 and 594 open reading frames (ORFs) and up to 29 tRNAs (Table 1). To measure the intrapopulation heterogeneity of pmoC-phages, we identified single nucleotide polymorphisms (SNPs) in BML_2, the most frequently detected pmoC-phage in BML samples, which showed a highly clonal population with little genetic diversity across different depths and sampling time points (Supplementary Fig. 13, Supplementary Information). No SNPs were detected in the pmoC gene indicating it is under purifying selection, while the genes with multiple SNPs included DNA polymerases and an endonuclease.

**Table 1.**
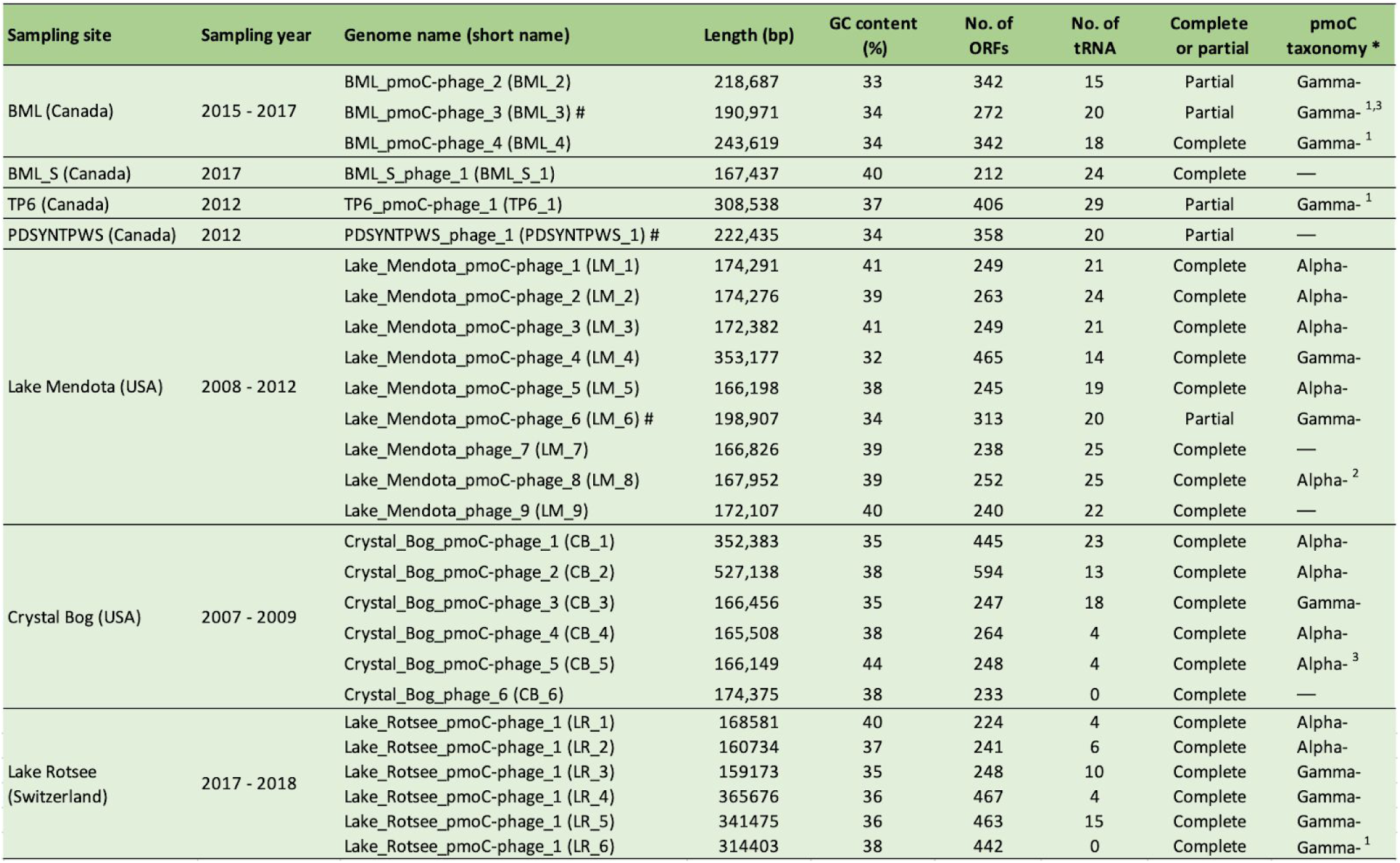
General features of the manually curated phage genomes. ^#^ Highly similar phage genomes, with identical large terminase and DNA polymerase sharing > 99.5% amino acid similarity. * The taxonomy is determined based on *pmoC* phylogeny including both phage and bacterial *pmoC* genes. ^1^ Fragmented *pmoC*, ^2^ partial *pmoC*, ^3^ some cells within the population only have partial *pmoC*. Note that no complete genome of pmoC-phage (though detected) was reconstructed from Trout Bog Lake samples due to the low sequencing coverage.

Notably, PDSYNTPWS_1 and BML_3 that were sampled from the same region of Canada but in different years share high genomic similarity with LM_6, but differ in the p*moC* region (Supplementary Fig. 14). PDSYNTPWS_1 does not contain the *pmoC* gene or the five neighboring genes found in BML_3 and LM_6 has *pmoC* but lacks the five neighboring genes. Interestingly, LM_1, LM_7 and LM_8 from Lake Mendota share a 2 kbp region near the *pmoC* of LM_8 that encodes hypothetical, phage-associated and bacterial genes, including part of an acyl-CoA dehydrogenase (Supplementary Fig. 15). This region present in these phages may be due to recombination that occurred during coinfection. The similarity of acyl-CoA dehydrogenase to a gene from *Methylocystis* spp. may indicate that this bacterium is the host (further discussed see below).

Eight published complete phage genomes (155-358 kbp in length) related to those reported here were retrieved based on viptree analyses (Supplementary Fig. 16) ^27,28^ and included in protein family analyses (Methods). Phylogenetic analyses based on the concatenated sequences of 11 universal phage specific proteins (Fig. 3, Supplementary Table 7) and DNA polymerases (Supplementary Fig. 17) suggested all pmoC-phages are *Myoviridae*. Generally, the similar the phage genome size the more closely phylogenetically related they are.

**Fig. 3.**
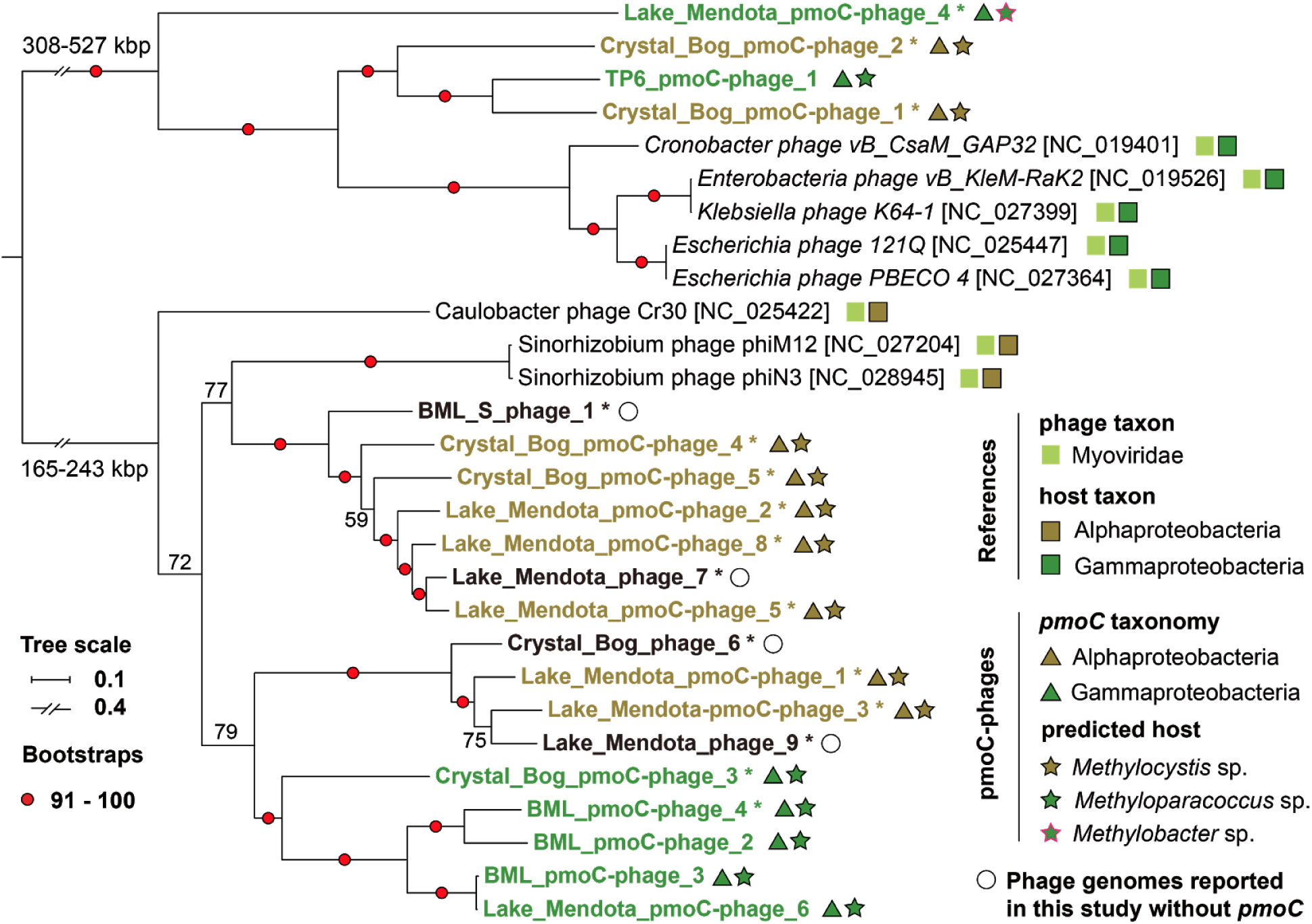
Phylogeny and predicted host of pmoC-phages. Phylogenetic analyses of pmoC-phages based on concatenated sequences of 11 universal proteins (encoded by single-copy genes) detected in the reconstructed phage genomes and the reference genomes (Supplementary Table 7). The 16 complete phage genomes reported in this study are indicated by asterisks, including four genomes of phages without *pmoC* (indicated by open circles). The genome size ranges of the two groups of phages are shown. The taxonomy of reference phages and their hosts are indicated by colored squares, and taxonomy of pmoC-phages and their predicted hosts by colored triangles and stars. The bootstraps are indicated by red circles when ≥ 91 or shown in numbers. See Supplementary Fig. 17 for phylogeny based on DNA polymerase sequences.

### Predicted hosts of pmoC-phages

CRISPR-Cas analyses found one spacer of Methyloparacoccus_57, and another spacer of a published *Methylobacter* genome, targeted the pmoC-phage BML_4 (Supplementary Fig. 18). However, none of the other pmoC-phage genomes were targeted by a CRISPR spacer from any coexisting microbes. Thus, we predicted their hosts using the similarity between the sequences of PmoC in phages and coexisting bacteria, assuming that *pmoC* was acquired by lateral transfer from their bacterial hosts ^3,6,29^ (Fig. 2b). Methyloparacoccus_57 was predicted as the host for the four oil sands pmoC-phages (including BML_3). The co-occurrence of Methyloparacoccus_57 and PDSYNTPWS_1, which is highly similar to BML_3, in the heavy PDSYNTPWS DNA-SIP fraction supports this. In LM samples, alphaproteobacterial and gammaproteobacterial methanotrophs were predicted as hosts of the pmoC-phages. One predicted host, *Methylocystis* sp. (an alphaproteobacterium), and the infecting pmoC-phages LM_1, LM_2, LM_3, LM_5 and LM_8 were detected together in all five years, especially in samples collected in Sep/Oct of each year (Supplementary Fig. 19). The phages LM_4 and LM_6 and their predicted gammaproteobacterial hosts (*Methylobacter* sp. and *Methyloparacoccus* sp., respectively) coexisted in the communities collected in 2012. The pmoC-phages from Crystal Bog were predicted to replicate in *Methylocystis* sp. (CB_1, _2, _4 and _5) and *Methyloparacoccus* sp. (CB_3) and time-series analyses verified that they coexisted in the communities (Supplementary Fig. 19). Together, these results strongly support the predicted host-phage relationships.

### Metabolic potentials of pmoC-phages and their relatives

We evaluated the protein families of pmoC-phages and related genomes to determine whether PmoC is associated with any other specific protein(s) (Fig. 4a, Supplementary Table 8). Generally, phylogenetically related phages have more similar protein families profiles. We found that PmoC is the only protein specific to all pmoC-phages (Fig. 4a). Genes for HSP20 were detected in all but two pmoC-phages and are encoded next to *pmoC* in five pmoC-phages. However, the significance of this is difficult to evaluate because HSP20 has been reported as a core gene of Cyanobacteria phages (Cyanophages) ^30^, which are phylogenetically related to the phages reported here (Supplementary Fig. 17). Moreover, all five related phages without *pmoC* also encode HSP20, suggesting that HSP20 is not related to the PmoC function (Fig. 4a).

**Fig. 4.**
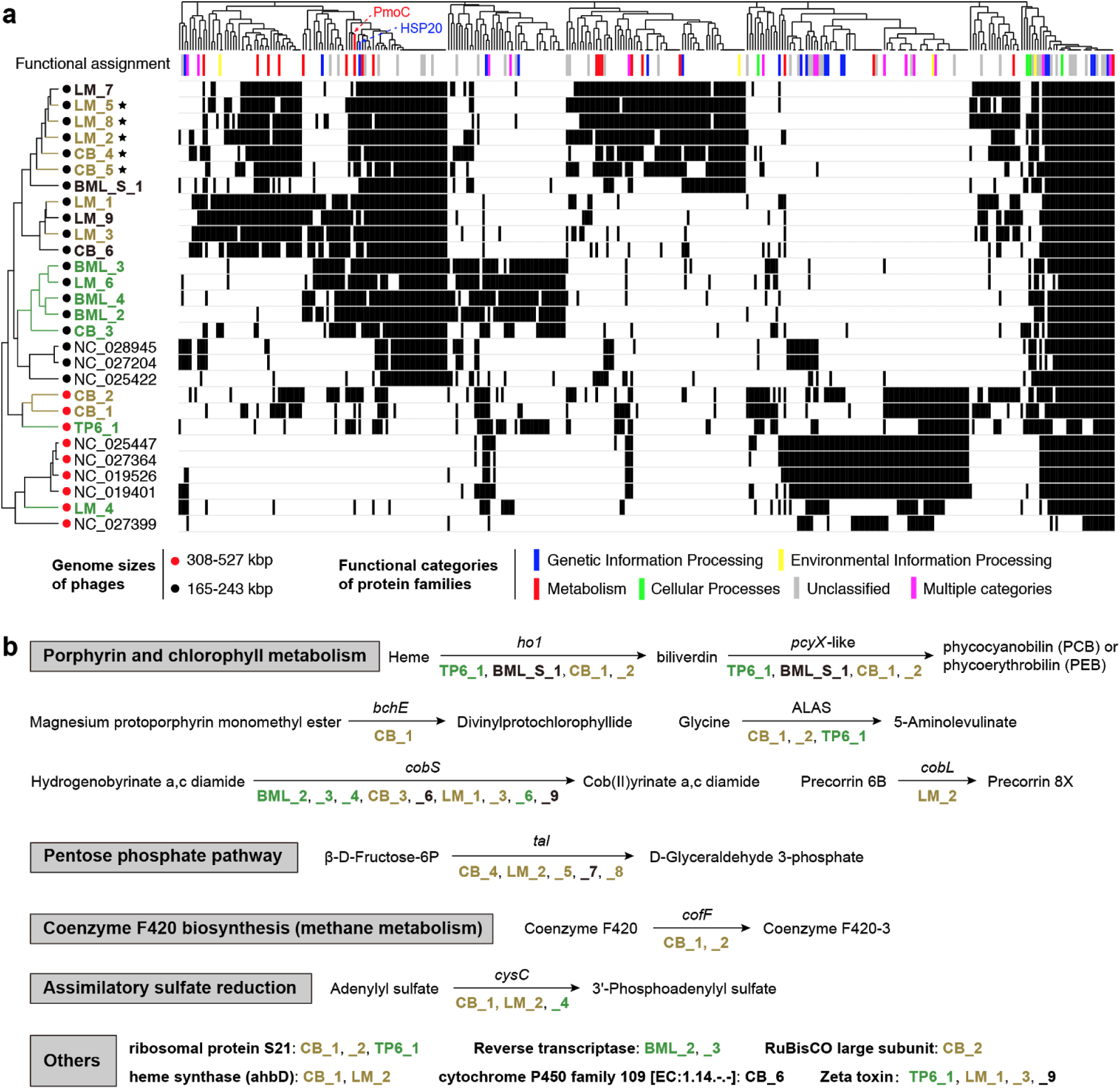
Metabolisms of pmoC-phages and their relatives. (a) Clustering of phages based on the presence/absence profiles of protein families that are encoded by at least five phages. The protein families were annotated using the kofam HMM database and assigned to functional categories, as indicated by colored bars (no bar for those without predicted function). The names of pmoC-phages infecting alpha- and gamma-proteobacterial methanotrophs are shown in gray and green respectively. The pmoC-phages with *pmoC* and HSP20 genes next to each other are indicated by stars. (b) Genes of specific metabolic potentials detected in newly reconstructed phage genomes. Abbreviations: *ho1*, heme oxygenase [EC:1.14.15.20]; *pcyX*-like, phycocyanobilin:ferredoxin oxidoreductase [EC:1.3.7.5]; *bchE*; anaerobic magnesium-protoporphyrin IX monomethyl ester cyclase [EC:1.21.98.3]; ALAS, 5-aminolevulinate synthase [EC:2.3.1.37]; *cobS*; cobaltochelatase CobS [EC:6.6.1.2]; *cobL*, precorrin-6Y C5,15-methyltransferase (decarboxylating) [EC:2.1.1.132]; *tal*, transaldolase [EC:2.2.1.2]; cofF; gamma-F420-2:alpha-L-glutamate ligase [EC:6.3.2.32]; cysC, adenylylsulfate kinase [EC:2.7.1.25].

Some other genes reported in Cyanophage genomes ^31^ were detected in phages reported here. Two genes found in both Cyanophages and Cyanobacteria from global ocean habitats ^32^, a heme oxygenase (*ho1*) and a *pcyX*-like phycocyanobilin:ferredoxin oxidoreductase are involved in the production of phycobiliprotein pigment ^32,33^. The *ho1* and *pcyX*-like genes are adjacent in TP6_1, BML_S_1 and CB_1 and one-gene-apart in CB_2 (Supplementary Fig. 20). One or more other genes for enzymes in porphyrin and chlorophyll metabolism (*bchE*, ALAS, *cobS, cobL*) were detected in TP6_1, CB_1, CB_2 and other related phages that lack *pmoC.* The transaldolase (*tal*) involved in the pentose phosphate pathway and reported in Cyanophages to enhance dNTP synthesis for phage replication ^29^, was encoded by five phages (4 are pmoC-phages).

Interestingly, *cofF* required for the biosynthesis of coenzyme F420 involved in methane metabolism is encoded by CB_1 and CB_2 pmoC-phages. Additionally, genes for assimilatory sulfate reduction and a zeta toxin (a bactericide that inhibits cell wall biosynthesis) were present in some phages reported here.

To investigate the gene expression of pmoC-phages *in situ*, we analyzed a set of metagenomic and metatranscriptomic datasets made public very recently ^34^, and obtained another 6 complete genomes of pmoC-phages that are phylogenetically related to those from our other study systems (Table 1, Supplementary Fig. 17). Transcriptional analyses showed high expression of the *pmoC* genes as well as genes encoding prohead and major capsid proteins (Supplementary Fig. 21). High *in situ* activities of phage-associated *pmoC* indicates the potential significance of these genes for phage during infection and supports our inference that phage-associated *pmoC* can impact overall rates of methane oxidation in freshwater ecosystems.

## Discussion

### PmoC-phages were overlooked in previous analyses

Previous cultivation-based studies isolated phages of bacterial methanotrophs from various habitats including oil waters and soil, but genomes of these phages have not been reported ^35,36^. Only 13 genome scaffolds of phages infecting methanotrophs (10-62 kbp in length) have been reported, and these sequences came from thawed permafrost samples ^37^. Here we described 16 large genomes of pmoC-phages that we propose can infect bacterial methanotrophs (up to 527 kbp; Table 1), but none of them are genomically or phylogenetically related to those from permafrost. Thus, bacterial methanotroph phages may be habitat specific. PmoC-phages have been overlooked in previous studies, in part because of focus on high level patterns such as global distribution, diversity and host specificity rather than gene inventories ^38^ and in part because of the high similarity between phage-associated and bacterial PmoC can fragment assemblies. The reconstruction of pmoC-phage genomes from multiple distinct habitats highlights the power of genome-resolved metagenomics and also the necessity of manual genome curation for accuracy ^39^.

### Why only *pmoC* in phages?

As analyzed in this study, *pmoC* but not *pmoA* and *pmoB* subunits of pMMO was detected in phages. Similarly, previous studies reported *amoC* was the only subunit of ammonia monooxygenase (homolog of pMMO) encoded by phages infecting Thaumarchaeota ^5,6^. Although possibly acquired from bacterial hosts along with other genes, only *pmoC* was retained as it can enhance phage fitness alone. In bacterial methanotrophs, there is increasing evidence indicating the essential role of PmoC in methane oxidation and the absolute necessity of PmoB is questionable ^12,25,26,40^. Given that the structure of PmoC is susceptible and largely disordered when the cell membrane is perturbed ^41^, we suggest that the additional pmoC genes either in the bacterial methanotroph or phage genome (Supplementary Table 3) could sustain methane oxidation under abnormal conditions. Further, the use of either Zn or Cu in the PmoC catalytic site raises the possibility that the additional PmoC provides resilience during metals limitation.

Previous studies suggested that different copies of *pmoC* ^12,42,43^ and *amoC* ^44^ from a single organism have distinct expression preferences under different conditions. Condition-dependent expression of *pmoC* is likely determined by sequence divergences in their termini, given that the middle regions are generally very similar (Supplementary Fig. 4). As the bacterial and phage-associated PmoC sequences are also only divergent in the N- and C- termini (Fig. 2a, Supplementary Fig. 9), the phage-associated PmoC may function as the standalone PmoC in bacterial methanotrophs under some conditions. Thus, termini divergences could increase the fitness of pmoC-phages following infection.

### Potential biogeochemical impacts of pmoC-phages

Anticorrelated abundance patterns of phage and predicted hosts suggest that pmoC-phages could reduce methane oxidation in an ecosystem by lysing their bacterial methanotroph hosts (Supplementary Fig. 19). On the other hand, pmoC-phages may accelerate methane oxidation during infection by providing PmoC variants. For example, Methyloparacoccus_57 was implicated as a major methane oxidizer in some BML samples and co-occurred with pmoC-phages that replicated recently in their bacterial hosts, given their relatively high abundance at the time of sampling (Fig. 1, Supplementary Information). Such effects may be important given that freshwater lake ecosystems are important sources of terrestrial methane emission ^15^

The presence of photosynthesis genes in pmoC-phage genomes is intriguing, given that they likely had to infect a cyanobacterial cell to acquire them. The pmoC-phages are phylogenetically related to Cyanophages, thus they may replicate in Cyanobacteria under conditions when they co-occur with them. The large genome size compared to most phages known to date may include genes required for host range expansion. Given that Cyanobacteria produce O_2_ that is required for methane oxidation by methanotrophs, and a very recent study indicated the production of methane by Cyanobacteria ^45^, it is possible that future work will show that pmoC-phages with broad host range can have far reaching impacts on the methane cycle.

## Conclusion

Our analyses based on genomic and geochemical information suggest the potential importance of pmoC-phages in methane oxidation and other biogeochemical cycling, and motivate the biochemical investigation of the role of phage-derived pmoC in the functioning of pMMO. The findings may be important to the understanding of methane emissions from freshwater ecosystems.

## Methods

### Sampling, DNA extraction and metagenomic analyses

The BML samples were collected from multiple depths of an end pit lake for oil sands wastes remediation in Alberta of Canada from 2015 to 2019 (Supplementary Table 1). The geochemical features of the samples were determined *in situ* or in the laboratory as previously described ^17^. Genomic DNA was collected filtering ca. 1.5 L water through 0.22-µm Rapid-Flow sterile disposable filters (Thermo Fisher Scientific) and stored at -20 °C until DNA extraction. DNA was extracted from the filters as previously described ^46^. The DNA samples were purified for library construction and sequenced on an Illumina HiSeq1500 platform with paired-end (PE) 150 bp kits. The LM and CB samples were collected from Lake Mendota (Madison, Wisconsin, USA) from 2008 to 2012 and Crystal Bog (Madison, Wisconsin, USA) from 2007 to 2009. The geochemical features and the procedures of sampling, DNA extraction and sequencing were detailed elsewhere ^47^, and the metagenomic reads were reassembled for pmoC-phages in this study. The raw reads of each metagenomic sample were filtered to remove Illumina adapters, PhiX and other contaminants with BBTools ^48^, and low-quality bases and reads using Sickle (version 1.33; https.github.com/najoshi/sickle). The high-quality reads of each sample were assembled using idba_ud ^49^ (parameters: --mink 20 --maxk 140 --step 20 --pre_correction). For a given sample, the high-quality reads of all samples from the same sampling site were individually mapped to the assembled scaffold set of each sample using bowtie2 with default parameters ^50^. The coverage of a given scaffold was calculated as the total number of bases mapped to it divided by its length. Multiple coverage values were obtained for each scaffold to reflect the representation of that scaffold in the various samples. For each sample, scaffolds with a minimum length of 3 kbp were assigned to preliminary draft genome bins using MetaBAT with default parameters ^51^, with both tetranucleotide frequencies (TNF) and coverage profiles of scaffolds considered. The scaffolds from the obtained bins and the unbinned scaffolds with a minimum length of 1 kbp were uploaded to the ggKbase platform. The genome bins of bacterial methanotrophs were evaluated based on the consistency of GC content, coverage and taxonomic information and scaffolds identified as contaminants were removed.

### Microbial community composition

The ribosomal protein S3 (rpS3) was used as a taxonomic marker gene for microbial community composition analyses. All the rpS3 proteins were predicted using hmmsearch ^52^ based on the tigrfam ^53^ HMM databases (TIGR01008 for Archaea and Eukaryotes, and TIGR01009 for Bacteria). The HMM hits were filtered by the tigrfam cutoff and searched against the NCBI RefSeq database ^54^ by BLASTp to remove those with the best hit of Eukaryotes. The retained bacterial and archaeal rpS3 amino acid sequences were clustered by cd-hit ^55^ with 100% similarity (-aL 0.8 -aS 0.8). The nucleotide sequences of all representative rpS3 were extracted and used as a dataset for reads mapping to calculate the coverage of them in each sample, which was performed by bowtie2 ^56^ with the default parameters. The coverage of a given scaffold was calculated only when the reads from a given sample could cover at least 50% of the nucleotide sequence. The relative abundance of a taxon in a given sample was calculated as the coverage of the corresponding rpS3 divided by the collective coverage of all representative rpS3 in the sample.

### Manual genome curation

The phage scaffolds were determined by phage specific genes as previously described ^57^, including capsid, phage, virus, prophage, terminase, prohead, tape measure, tail, head, portal, DNA packaging. The confirmed phage scaffolds were predicted for protein-coding genes using Prodigal ^58^ and searched against the HMM databases of proteins involved in methane metabolisms. The phage scaffolds with *pmoC* genes and also a minimum sequencing coverage of 20X were manually curated to completion and/or to fix any assembly errors following the pipelines as described previously ^39^. Manual fixation of assembled errors and extension to completion of phage genomes are time-consuming but essential to reveal their metabolic potentials. In detail, firstly an overlap-based assembly of scaffolds was performed using Geneious ^59^, then linkage of scaffolds by scaffold extension, and manual fixation of local assembly errors detected by ra2.py ^60^. All the curated phage genomes were manually checked and confirmed by mapping the reads to the final genome for accuracy. The pmoC-phage scaffold of TP6_1 was sequenced by 454 pyrosequencing and Illumina ^21^ and was extended by overlap at the ends of scaffolds detected by BLASTn using 454 reads followed by confirmation of the extension by Illumina reads. The BLASTn search and extension were performed several times until no more scaffolds with end overlap could be found. For the genomes of phages closely related to pmoC-phages, we firstly identified the scaffolds by searching against all the large terminase (TerL) proteins from pmoC-phages, and those scaffolds having a TerL with ≥ 80% amino acid similarity were selected as targets for scaffold extension and genome curation to completion. The phylogenetic and protein family analyses supported our determination of related phages based on TerL similarity. The similarity of phage genomes was calculated using the online average nucleotide identity tool ^61^. Attempt to retrieve highly similar phage genomes from published NCBI SRA datasets was performed using the online tool ^62^. For genomes of bacterial methanotrophs, the local assembly errors were fixed as performed for phage genomes. For the pMMO operon and standalone *pmoC* scaffolds of the bacterial methanotrophs, we generally manually curated them using the reads from the sample(s) without pmoC-phages detected. A total of 51 bacterial universal single-copy genes (SCGs) were used to evaluate genome completeness and contaminations ^63^.

### Bacterial sMMO and pMMO subunits

To reveal the sMMO and pMMO subunits in published genomes of bacterial methanotrophs, all the genomes assigned to the well-known methanotroph genera ^64^ were downloaded from NCBI (Supplementary Table 3), along with their protein sequences and annotation information. The standalone *pmoC* genomes in published genomes were determined manually based on their genomic context. For the bacterial methanotrophs with genomes reconstructed in this study, their protein-coding genes were predicted using Prodigal ^58^ and searched against the sMMO and pMMO subunits HMM databases from TIGRFAM ^53^. The corresponding scaffolds were manually checked for potential assembly errors and fixed once found.

### CRISPR-Cas analyses

All the predicted proteins of scaffolds with a minimum length of 1 kbp were searched against local HMM databases including all reported Cas proteins, and the nucleotide sequences of the same set of scaffolds were scanned for CRISPR loci using minced ^65^ (-minSL = 17). The spacers were extracted from the scaffolds with CRISPR loci as determined by minced, and the reads that could be mapped to the scaffolds with a local python script as previously described ^57^(only spacers from the scaffolds of published methanotroph genomes were extracted, as no mapped reads are available). Duplicated spacers were removed using cd-hit-est and the unique ones were built as a database for BLASTn search (task = blastn-short; e-value = 1e-3) with the pmoC-phage scaffolds from the same sampling site as queries. Once a spacer was found to target a pmoC-phage scaffold (≥ 30 bp), the original scaffold of the spacer was manually confirmed for CRISPR locus and Cas proteins.

### Distribution of phages and their predicted hosts

The quality reads from each sampling site were mapped to the genomes of pmoC-phages reconstructed from the same site. The occurrence of a given phage in a given sample was determined if ≥ 75% of its genome could be covered by reads with ≥ 95% nucleotide similarity. The occurrence of a given predicted host was determined if its genome (or scaffold containing the pmoC that most similar to the one in pmoC-phage) was mapped by reads with ≥ 98% nucleotide similarity (and when ≥ 75% of the genome or scaffold was covered). The sequencing coverage of a given pmoC-phage or predicted host genome/scaffold in a sample was calculated by the total length of mapped reads dividing by the length of the corresponding genome/scaffold. See Supplementary Information for details of how to determine the pmoC-phages and their predicted host in published oil sands related metagenomic datasets. Methyloparacoccus_57 could be detected in Lake Mendota samples (with identical 16S rRNA gene sequence found) but with very low sequencing coverage, thus no quality genome is obtained. Given the high similarity between LM_6 and BML_3, we predicted that Methyloparacoccus_57 was the host of LM_6, and the genome of Methyloparacoccus_57 from BML was used to profile its existence across the Lake Mendota samples as described above.

### Phage protein family analyses

Protein family analyses were performed as previously described ^66^. In detail, first, all-vs-all searches were performed using MMseqs2 ^67^, with parameters set as e-value = 0.001, sensitivity = 7.5 and cover = 0.5. Second, a sequence similarity network was built based on the pairwise similarities, then the greedy set cover algorithm from MMseqs2 was performed to define protein subclusters (i.e., protein subfamilies). Third, in order to test for distant homology, we grouped subfamilies into protein families using an HMM-HMM comparison procedure as follows. The proteins of each subfamily with at least two protein members were aligned using the result2msa parameter of MMseqs2, and HMM profiles were built from the multiple sequence alignment using the HHpred suite ^68^. The subfamilies were then compared to each other using hhblits ^69^ from the HHpred suite (with parameters -v 0 -p 50 -z 4 -Z 32000 -B 0 -b 0). For subfamilies with probability scores of ≥ 95% and coverage ≥ 0.5, a similarity score (probability × coverage) was used as the weights of the input network in the final clustering using the Markov CLustering algorithm ^70^, with 2.0 as the inflation parameter. Finally, the resulting clusters were defined as protein families. The clustering analyses of the presence and absence of protein families detected in the phage genomes were performed with Jaccard distance and complete linkage.

### Phylogenetic analyses

Phylogenetic analyses were performed for bacterial and phage-associated PmoC sequences identified from BML, BML_S, LM, CB and TBL samples, with NCBI bacterial methanotrophs PmoC protein sequences (see above) included for references. To reveal the phylogeny of phages with genomes reconstructed in this study, sequences of 11 protein subfamilies retrieved from the protein family analyses (see above) were concatenated for analyses. In addition, the DNA polymerase (within the 11 proteins used for concatenation) was used as a single marker for phylogenetic analyses. All DNA polymerase of NCBI RefSeq viruses/phages were downloaded and used to retrieve references by BLASTp (using the DNA polymerase sequences reported in this study as queries). The top 30 BLASTp hits were included as references.

For the phylogeny of bacterial methanotrophs, 16 concatenated ribosomal proteins (16RPs) ^71^, ribosomal protein S3 (rpS3) and 16S rRNA gene sequences were used as markers. For protein-coding genes predicted by prodigal ^58^ from scaffolds with a minimum length of 1 kbp, the 16 RPs (including rpS3) were determined by HMM-based search with databases built from Hug et al. ^71^. For those scaffolds with 8 or more of the 16RPs, the ribosomal proteins were individually aligned and filtered. Another tree only based on rpS3 was constructed using the same procedure. The references for both 16RP and rpS3 trees were selected from the Hug et al. ^71^ datasets by rpS3 BLASTp search with the top 5 hits included. The 16S rRNA genes were predicted via HMM search as previously described ^60^, and any insertion with a minimum length of 10 bp was removed. The insertion-removed 16S datasets were aligned using a local version of SINA aligner ^72^ and filtered by trimAl to remove those columns with ≥ 90% gaps. The tree was built by IQtree ^73^ using the “GTR+G4” model. References were selected based on BLASTn search against the 16S datasets of Silva132 ^74^, and the top 5 hits were included. For all the phylogenetic analyses with protein sequences, the proteins were aligned using Muscle ^75^ and filtered by trimAl ^76^ to remove those columns with ≥ 90% gaps, followed by tree building with IQtree ^73^ using the “LG+G4” model, filtered sequences were concatenated for multiple proteins based analyses.

## Data availability

The genome of Methyloparacoccus_57 that reconstructed from the BML datasets has been deposited at NCBI under BioProject xx (TBA), which is also available at Figshare (TBA). All the phage genomes and are available through Cyverse at https://de.cyverse.org/dl/d/85779809-E882-4C58-9E7B-07912799AFB3/my.final.20.phages.fasta.

## Author contributions

L.X.C. designed the analyses. T.C.N. collected and prepared the BML and BML_S samples for sequencing. T.C.N. and L.A.W. provided the DNA sequencing of BML and BML_S samples. K.D.M. provided the metagenomic datasets of LM, CB and TBL samples. L.X.C. performed the metagenomic assembly, genome binning, genome annotation, phylogenetic analyses, HMM search, and CRISPR-Cas analyses. L.X.C. and J.F.B. performed manual genome curation to completion. R.M. and L.X.C. performed protein family analyses. A.C.C. performed the SNPs analysis. L.X.C. and J.F.B. wrote the manuscript. All authors read and approved the final manuscript.

## Acknowledgements

We thank Dr. Magdalena J. Mayr for the permission to use the metagenomic and metatranscriptomic datasets from Lake Rotsee for analyses in this study. We thank Dr. Robert Edwards’s help in attempting to retrieve highly similar phage genomes in NCBI SRA datasets. The study was supported by the NSERC Canada and Syncrude Canada (Grant No. CRDPJ 403361–10). We also acknowledge funding support from the Chan Zuckerberg Biohub and the Innovative Genomics Institute at UC Berkeley. Katherine D. McMahon received funding from the United States National Science Foundation Microbial Observatories program (MCB-0702395), the Long-Term Ecological Research Program (NTL–LTER DEB-1440297), and an INSPIRE award (DEB-1344254).

