## Supplementary Info and Figs for "Large Freshwater Phages with the Potential to Augment Aerobic Methane Oxidation"

**Supplementary information**
**for**

### Large Freshwater Phages with the Potential to Augment Aerobic Methane Oxidation

Lin-Xing Chen[1], Raphaël Méheust[1], Alexander Crits-Christoph[2], Katherine D. McMahon[3], Tara Colenbrander Nelson[4], Lesley A. Warren[4,5], and Jillian F. Banfield[1,2,6,7]*

[1] Department of Earth and Planetary Sciences, Berkeley, CA, USA.
[2] Department of Plant and Microbial Biology, University of California, Berkeley, Berkeley, CA, USA
[3] Departments of Civil and Environmental Engineering, and Bacteriology, University of Wisconsin, Madison, WI 53705, USA
[4] Department of Civil and Mineral Engineering, University of Toronto, Toronto, Canada
[5] School of Geography and Earth Science, McMaster University, Hamilton, Canada
[6] Department of Environmental Science, Policy, and Management, Berkeley, CA, USA.
[7] Earth and Environmental Sciences, Lawrence Berkeley National Laboratory, Berkeley, CA, USA.

*Corresponding author:

Address: McCone Hall, Berkeley, CA 94720

**Running title: Phages that augment methane oxidation**

**Re-analyses of published oil sands datasets**

Datasets from four previously published studies related to oil sands waste lakes are included for analyses here, see below for details. To sum, studies 1 and 2 were detected with both Methyloparacoccus_57 and pmoC-phages, studies 3 and 4 with only Methyloparacoccus_57.

***1. Tan et al., 2015 - Comparative analysis of metagenomes from three methanogenic hydrocarbon-degrading enrichment cultures with 41 environmental samples.***

We firstly analyzed the datasets of enrichment samples added with short-chain alkane (C6-C10; SCADC) or naphtha (NAPDC) or toluene (TOLDC), and did not detect Methyloparacoccus_57 or any pmoC-phages. Then we analyzed the other two metagenomic datasets used for comparison in the original paper, i.e., TP6 and TP_MLSB.

The sample TP6 (UTM 466358E 6319838N) was collected in 2012 from Suncor tailing pond at the depth of 6 meters and sequenced with both 454 pyrosequencing and Illumina (accession number: SRX327722). We detected one pmoC-phage (referred to as "TP6_1") from the original assembly and extended the TP6_1 genome using the 454 pyrosequencing reads and Illumina to the current version (see Table 1 and descriptions in the main text). No other pmoC-phage identified in BML samples was detected in this sample. For its host, we compared the PmoC sequence of TP6_1 to all others from the assembly, and analyzed all the bacterial and archaeal species in the community via rpS3 phylogeny for methanotroph(s), and found that the host of TP6_1 is Methyloparacoccus_57 that reported in this study (see main text).

The sample TP_MLSB was collected from Syncrude in 2011 (accession number: SRR636569), the quality Illumina reads were downloaded and mapped to genomes reconstructed from BML (with > 98% nucleotide identity). As a result, Methyloparacoccus_57 (sequencing coverage: 7.37 X, genome covered: 97.8%) and pmoC-phages of TP6_1 (sequencing coverage: 5.24 X, genome covered: 97.8%) and BML_3 (sequencing coverage: 7.01 X, genome covered: 89.6%) were detected. (Extended Data Fig. 1). We thus did not assemble this dataset because of the low sequencing coverage of these genomes. Interestingly, the *pmoC* region of BML_3 was only mapped by two reads, indicating that the corresponding phage in TP_MLSB generally did not contain *pmoC*. Also, the coverage peaks may indicate the existence of other related phage(s) and/or repeat regions in the genome of the same phage.

*In summary, by re-analyzing the datasets reported by Tan et al. 2015, we found one pmoC-phage (i.e., TP6_1) in the sample of TP6 from a Suncor tailing pond, and we showed the existence of phages similar to TP6_1 and BML_3 in the sample of TP_MLSB collected from Syncrude, and we detected Methyloparacoccus_57 in both samples.*

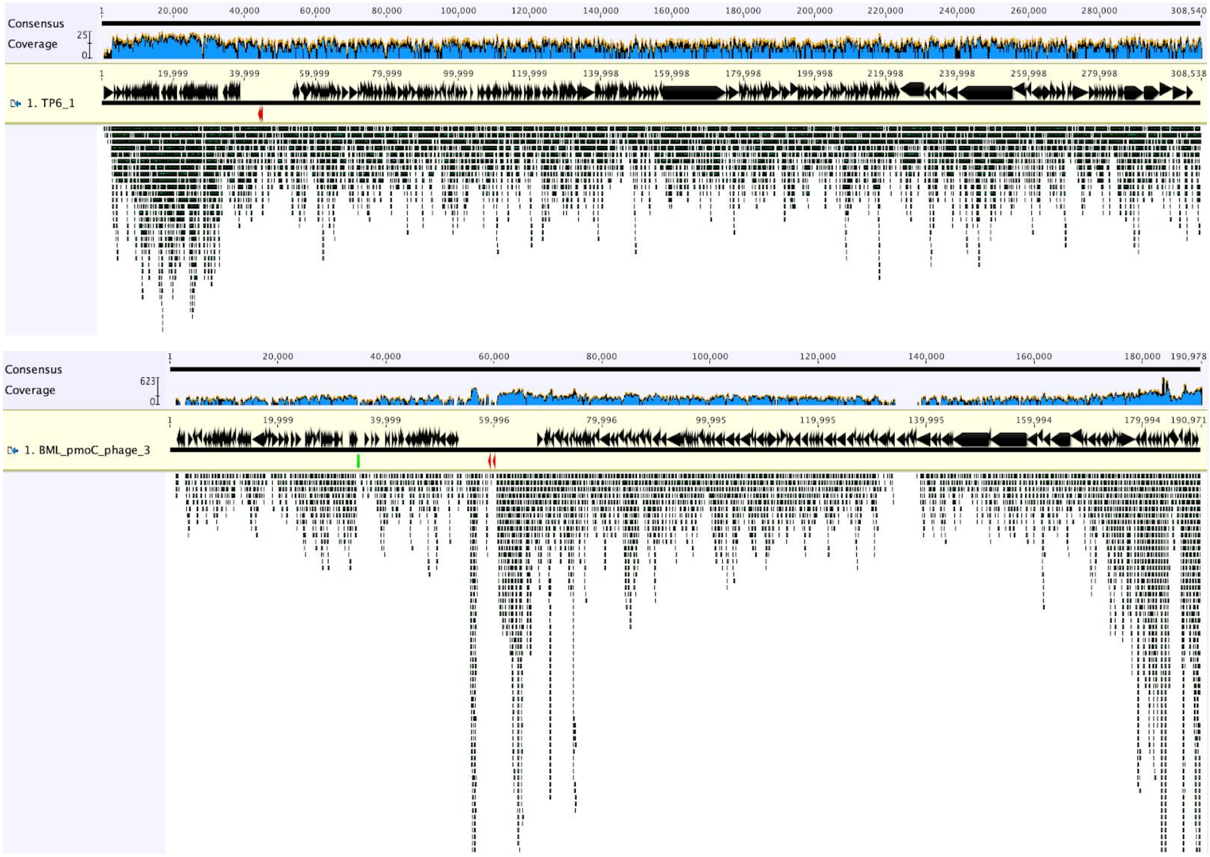


Extended Data Fig. 1 | Screenshot showing the mapping of reads from TP_MLSB to pmoC-phage genomes of TP6_1 (upper panel) and BML_3 (bottom panel). The *pmoC* genes are shown in red.

***2. Saidi-Mehrabad et al., 2013 - Methanotrophic bacteria in oilsands tailings ponds of northern Alberta.***

Saidi-Mehrabad et al. collected surface water (0–10 cm) at 1–3-month intervals over 2010–2011 from two tailings ponds near Fort McMurray, Alberta, Canada (i.e., Pond A and Pond B as designated in the original paper). As described in the original paper, "An aerobic methanotroph belonging to the Methylococcus/Methylocaldum cluster of Gammaproteobacteria (OTU12103) was among the predominantly detected OTUs in Pond A, making up on average 1.5% of all reads", we thus performed *de novo* assembly of the metagenomic dataset of PD_SYN_TP_WS_002_003_071511 (accession number: SRX327520; referred to as "PDSYNTPWS" hereafter) sequenced by Illumina and found that the predominant OTU12103 should be Methyloparacoccus_57 that reported in this study, as the 16S rRNA gene sequence from the assembly shared 100% similarity with that of Methyloparacoccus_57. Phylogenetic and sequencing coverage analyses also indicated Methyloparacoccus_57 (or related species) is the most abundant bacterial methanotroph in the community (Extended Data Fig. 2). Binning and subsequent curation obtained the Methyloparacoccus_57 related genome from PDSYNTPWS, which was referred to as "Methyloparacoccus_57_PDSYNTPWS".

We also mapped the Illumina reads to the pmoC-phage genomes reconstructed from BML and TP6 (see above), and found that the existence of phages similar to BML_3 and BML_4 (Extended Data Fig. 3), which was confirmed by BLASTn of assembled scaffolds against BML_3 and BML_4. Attempt for complete genome of phage similar to BML_3 (because it had sufficient sequencing coverage) obtained a high-quality genome (referred to as "PDSYNTPWS_1") (Extended Data Fig. 4). The genomic alignments of PDSYNTPWS_1, BML_3 and LM_6 are shown in Supplementary Fig. x and described in the main text.

In addition, DNA stable isotope probing (SIP) analyses using $^{13}$CH4 were conducted to track the active methane oxidizers in the PDSYNTPWS sample (Saidi-Mehrabad et al. 2013). A "Five microliters of a selected 'heavy' SIP fraction" of DNA sample was collected for amplification and sequencing for metagenomic analyses. The resulting Illumina reads (382 million read pairs) were downloaded for re-analyses here, including mapping to the genomes of Methyloparacoccus_57 and pmoC-phages reconstructed from BML in this study. As a result, ~ 6.58% of the metagenomic DNA-SIP reads could be mapped to Methyloparacoccus_57 (Extended Data Fig. 5), and some reads were mapped to the phage genome of PDSYNTPWS_1 (Extended Data Fig. 6). The uneven depth across the scaffold and the genome may be due to the multiple displacement amplification (MDA) in DNA preparation. We also performed *de novo* assembly of the DNA-SIP data and obtained a total length of 90 Mbp scaffolds, phylogenetic analyses based on rpS3 indicated that Methyloparacoccus_57 and some other gammaproteobacterial methanotrophs in the community were active in methane oxidation (Extended Data Fig. 7).

Saidi-Mehrabad et al. also reported a total of 22 16S rRNA gene datasets (sequenced by 454 GS FLX Titanium) in the original paper, while only 17 of them could be downloaded from NCBI SRA via the accession number provided (SRP013946). The 16S rRNA gene sequences were searched against that of Methyloparacoccus_57 reported in this study using BLASTn (> 98% similarity, > 500 alignment length), the total number of hits and the calculated relative abundance are shown in Extended Data Fig. 8 for each sample. As we could not match the NCBI SRA datasets to the samples described in the original paper, we thus show both the SRA accession number and sample description in the figure.

*In summary, by re-analyzing the datasets reported by Saidi-Mehrabad et al. 2013, Methyloparacoccus_57 were detected in most of the samples, we also obtained the genome of phage (i.e., PDSYNTPWS_1) highly similar to BML_3 while without pmoC. The DNA-SIP data showed Methyloparacoccus_57 is the predominant species for methane oxidation and PDSYNTPWS_1 also took the $^{13}$C for synthesis, indicating their potential host/phage relationship as described in the main text.*

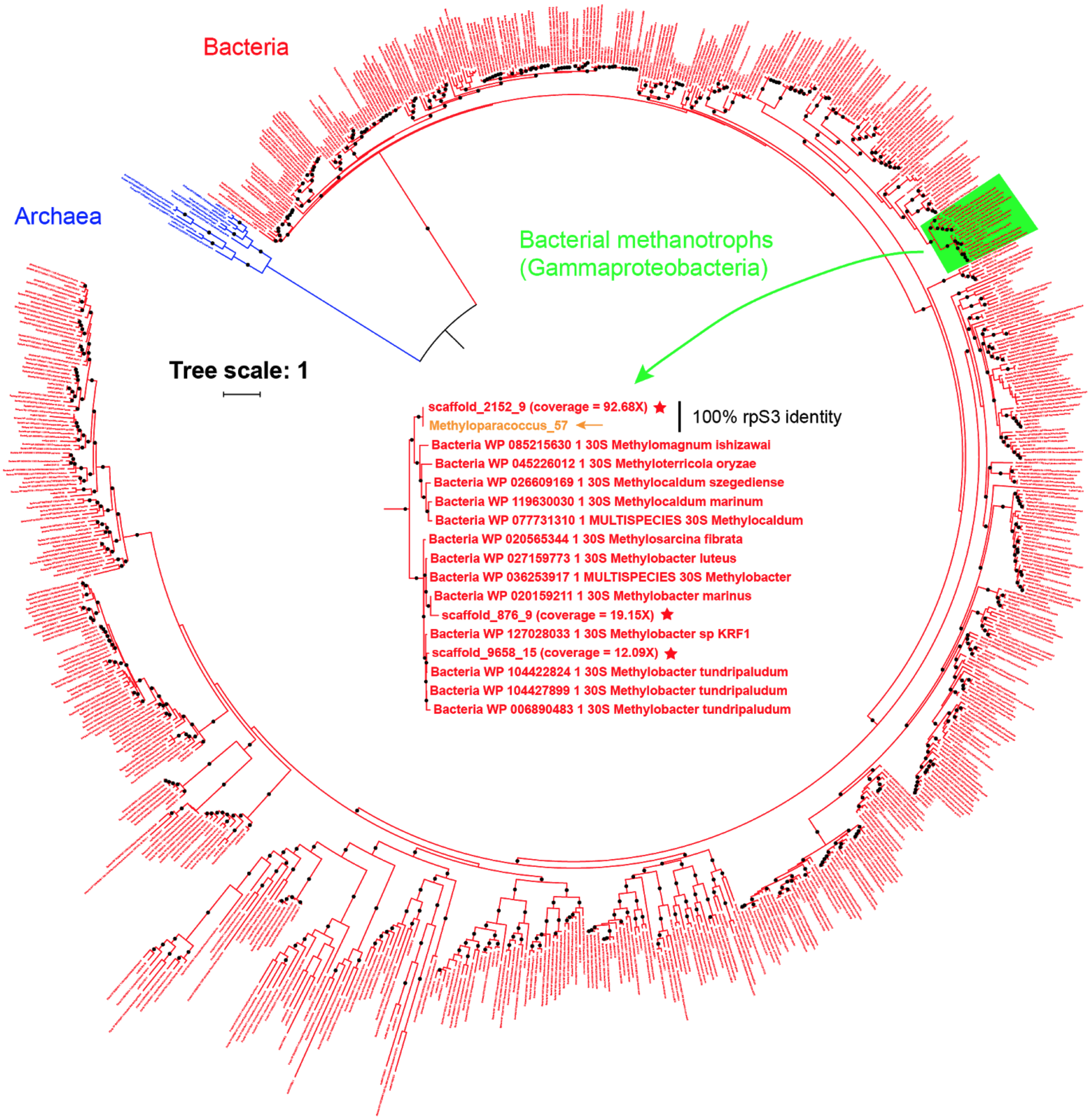


Extended Data Fig. 2 | Phylogenetic analyses based on rpS3 showing the bacterial methanotrophs (only gamma-) detected in the PDSYNTPWS sample. The phylogeny of the methanotrophs is zoomed-in in the middle, those from PDSYNTPWS are indicated by red stars and their sequencing coverages of the corresponding scaffolds are listed in the brackets, the rpS3 of Methyloparacoccus_57 (from BML) is included for reference. A black solid circle indicates bootstrap ≥ 70.

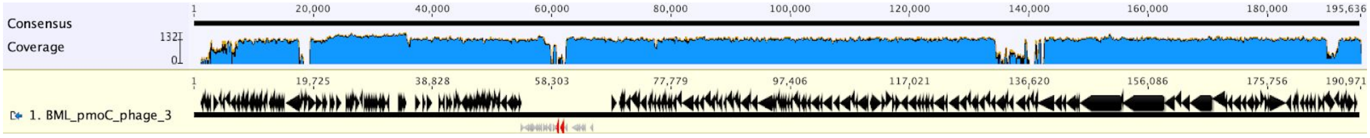


↑BML_3 (sequencing coverage ~ 40X)

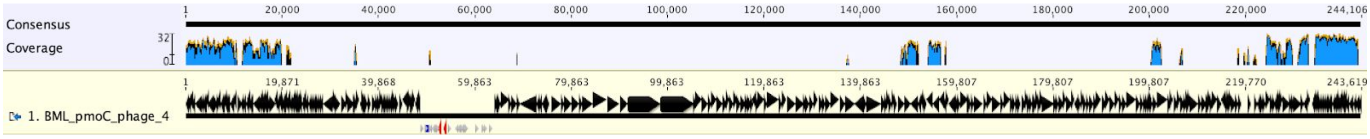


↑BML_4 (sequencing coverage ~ 10X)

Extended Data Fig. 3 | The detection of DNA-SIP metagenomic reads (highlighted in the red boxes) mapped to the genomes of pmoC-phages of BML_2, BML_3 and BML_4 indicating the existence of related phages in the sample. The mapping was performed by bowtie2 and filtered with no more than 2 mismatches. The *pmoC* genes are shown in red.

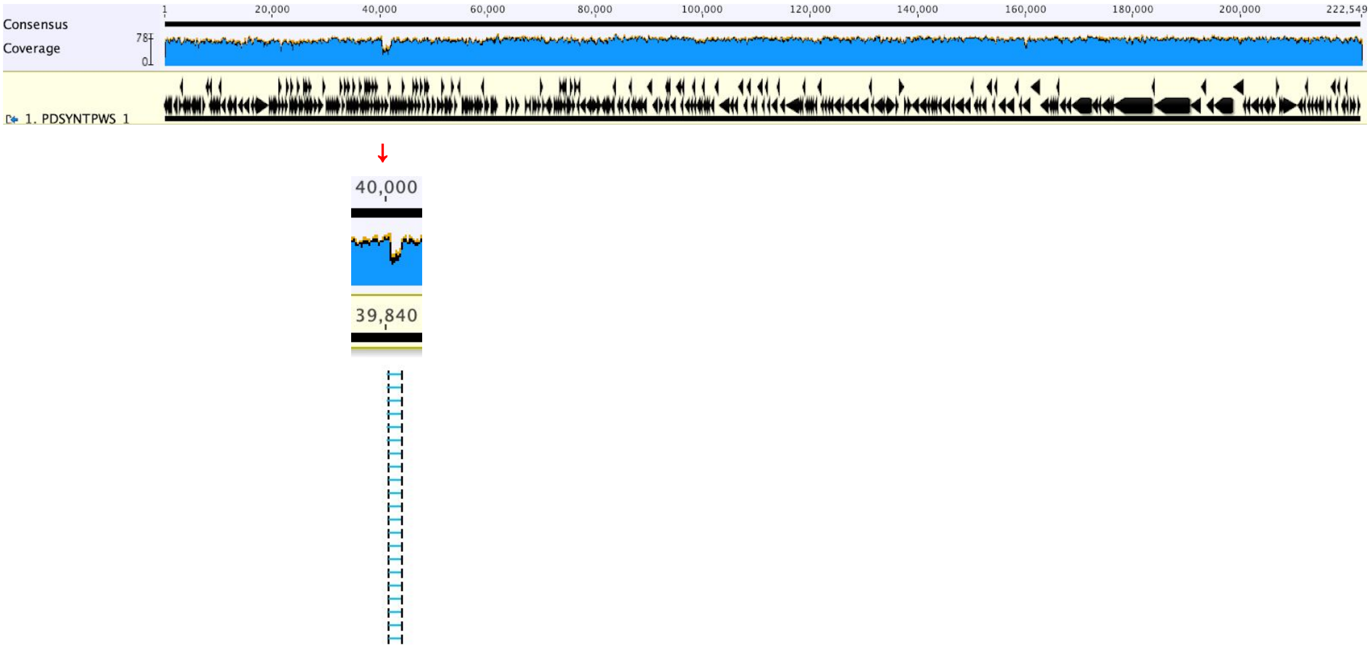


Extended Data Fig. 4 | The high quality of the phage genome of PDSYNTPWS_1 (highly similar to BML_3), reconstructed from the metagenomic dataset reported by Saidi-Mehrabad et al. 2013. The region with lower coverage is due to its absence in some of the phages, as indicated by the spanned read pairs.

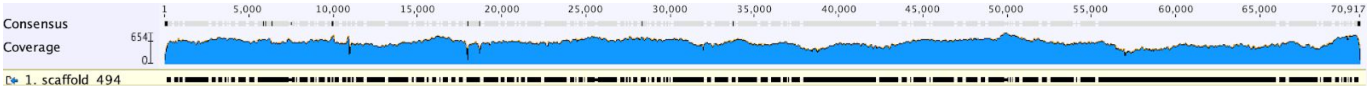


Extended Data Fig. 5 | Reads mapping (≥ 98 nucleotide identity) to scaffold of Methyloparacoccus_57_PDSYNTPWS in the heavy DNA-SIP sample. The longest scaffold of Methyloparacoccus_57_PDSYNTPWS (scaffold_494; 70,351 bp) is used as an example and shown here.

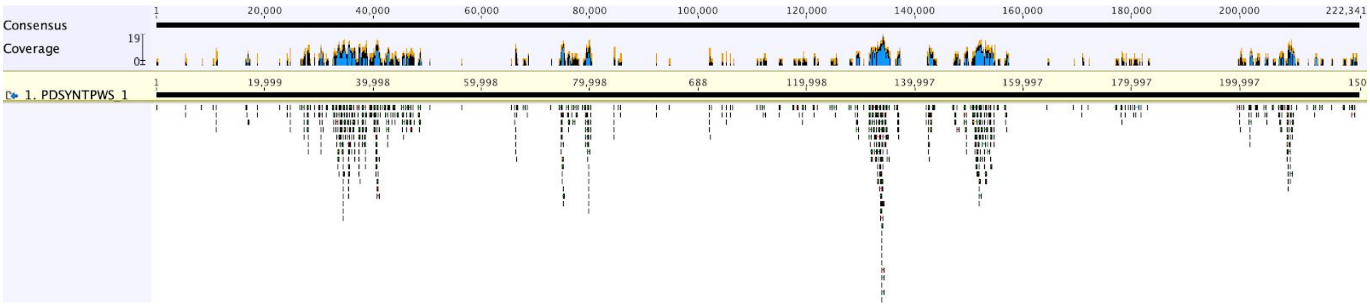


Extended Data Fig. 6 | Mapping of reads showing the usage of $^{13}CH_4$ by PDSYNTPWS_1 in the community analyzed by DNA-SIP. Reads mapping was performed by Bowtie2 and filtered allowing with > 98% nucleotide identity. The uneven depth may be due to the multiple displacement amplification in sequencing sample preparation (see Saidi-Mehrabad et al. 2013 for detail).

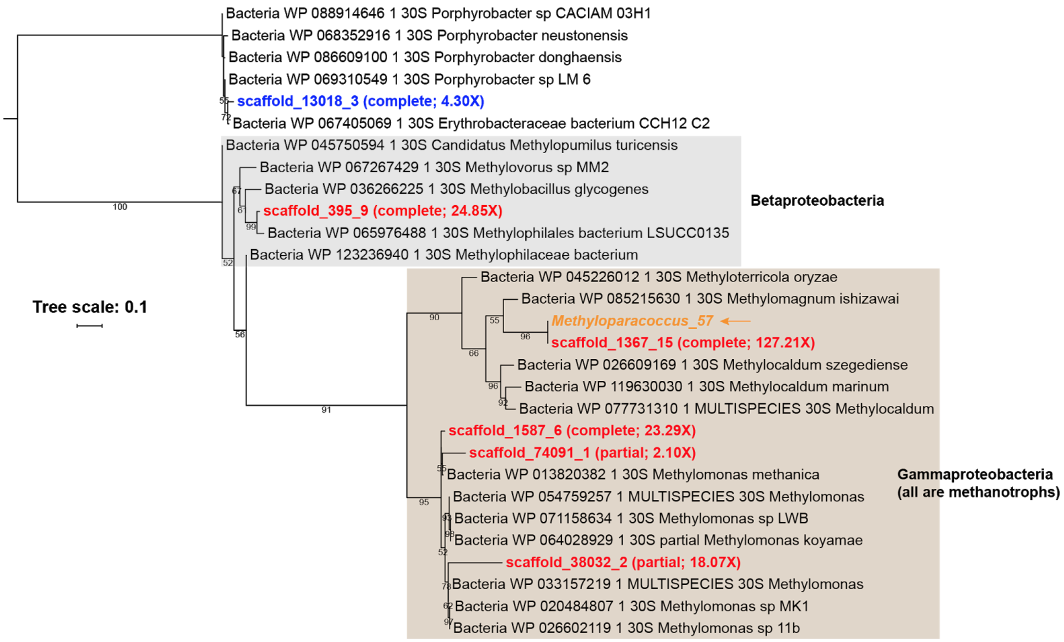


Extended Data Fig. 7 | Phylogenetic analyses showing the active members in the community as revealed by DNA-SIP analyses. The phylogeny was performed based on the rpS3 protein sequences, rpS3 of Methyloparacoccus_57 (indicated by an arrow) was included for reference. The sequencing coverages of the corresponding scaffolds are shown in the brackets.

| SRA accession ID | SRA description | # Total 16S reads | # Methylococcaceae_57 reads | Relative abundance |
|---|---|---|---|---|
| SRR516405 | PD_SUN_TP_WS_002_006_092509 | 13297 | 0 | 0.00% |
| SRR516408 | PD_SUN_OS_WS_001_002_092509 | 14901 | 0 | 0.00% |
| SRR516409 | PD_SUN_OS_WS_002_002_092509 | 5575 | 0 | 0.00% |
| SRR516411 | PD_SUN_TP_WS_001_006_092509 | 18846 | 14 | 0.07% |
| SRR516412 | PD_SUN_TP_WS_003_006_092509 | 14414 | 11 | 0.08% |
| SRR516413 | PD_SUN_TP_WS_004_006_092509 | 17616 | 15 | 0.09% |
| SRR516414 | PD_SUN_TP_WS_005_006_092509 | 28042 | 26 | 0.09% |
| SRR516415 | PD_SYN_TP_WS_001_006_072210 | 6386 | 2 | 0.03% |
| SRR516416 | PD_SYN_TP_WS_001_002_081211 | 16196 | 4 | 0.02% |
| **SRR516417** | **PD_SYN_TP_WS_002_003_071511** | **17125** | **79** | **0.46%** ← with Illumina metagenomic reads and also analyzed by DNA-SIP in lab. |
| SRR516420 | PD_SYN_TP_WS_003_003_071511 | 8257 | 17 | 0.21% |
| SRR516421 | PD_SYN_TP_WS_002_002_090111 | 17898 | 91 | 0.51% |
| SRR516422 | PD_SYN_TP_WS_002_002_081211 | 14344 | 66 | 0.46% |
| SRR516423 | PD_SYN_TP_WS_002_006_072210 | 7721 | 1 | 0.01% |
| SRR516424 | PD_SYN_TP_WS_001_003_071511 | 17557 | 3 | 0.02% |
| SRR516425 | PD_SYN_TP_WS_001_002_090111 | 19197 | 1 | 0.01% |

Extended Data Fig. 8 | The information of Methyloparacoccus_57 detected in the 16S rRNA gene sequence datasets reported by Saidi-Mehrabad et al., 2013.

***3. An et al., 2013 - Metagenomics of hydrocarbon resource environments indicates aerobic taxa and genes to be unexpectedly common.***

A total of 12 metagenomic datasets (sequenced by 454 pyrosequencing or Illumina) from oil sands related habitats were reported in this study. Methyloparacoccus_57 were only detected in PDSYNTPWS (454 pyrosequencing reads) by 16S rRNA gene sequence search, and genomic fragments similar to PDSYNTPWS_1 as well in the sample (Extended Data Fig. 9). Samples from the same site were also reported in **Saidi-Mehrabad et al., 2013** (see above). No pmoC-phage was detected in other samples reported in An et al., 2013.

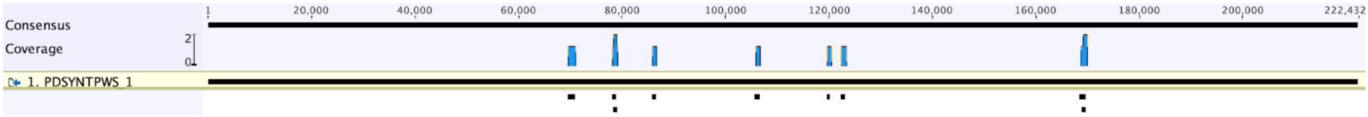


Extended Data Fig. 9 | Screenshots showing the alignment of 454 pyrosequencing reads from PDSYNTPWS in An et al., 2013, to the genome of PDSYNTPWS_1 reconstructed from **Saidi-Mehrabad et al., 2013**. The limited number of reads aligned was likely due to the low sequencing coverage of 454 pyrosequencing, and/or the low abundance of related phage in the sample, or genetic divergences (only reads with > 97% nucleotide identity were included for alignment).

***4. Rochman et al., 2017 - Benzene and Naphthalene Degrading Bacterial Communities in an Oil Sands Tailings Pond.***

Oil sands process-affected water (OSPW) was collected in 2012 for incubation experiment with the addition of Benzene and Naphthalene, to reveal the microorganisms in OSPW for their degradation, one control water sample was also analyzed via 16S analyses (sequenced by 454 GS FLX Titanium). The 16S rRNA gene sequence datasets were downloaded from NCBI SRA via the accession number of SRP109130 provided in the original paper, and compared against that of Methyloparacoccus_57 reported in this study by BLASTn (> 98% similarity, > 500 alignment length), the results are shown in Extended Data Fig. 10. As a result, the analyses indicate that Methyloparacoccus_57 was not the primary player in using methanol, naphthalene or benzene, however, it was with high abundance in the natural OSPW samples.

| SRA accession ID | SRA description (description in the paper) | # Total 16S reads | # Methylococcaceae_57 reads | Relative abundance |
|---|---|---|---|---|
| SRR5681085 | Methanol Heavy SIP | 9961 | 18 | 0.181% |
| SRR5681086 | Naphthalene Heavy SIP (Naphthalene) | 9705 | 6 | 0.062% |
| SRR5681088 | Benzene Heavy SIP (Benzene) | 13101 | 1 | 0.008% |
| SRR5681087 | Control (OSPW) | 712899 | 41003 | 5.752% |
| SRR5681089 | Heavy SIP (OSPW heavy) | 7111 | 248 | 3.488% |

Extended Data Fig. 10 | The information of Methyloparacoccus_57 detected in the 16S rRNA gene sequence datasets reported by Rochman et al., 2017.

**SNPs analyses of pmoC-phages**

As a case study, we investigated the population heterogeneity of the most commonly observed pmoC-phage in BML, BML_2, which was detected in 13 samples with ≥ 5X coverage.

We found that the genomes were remarkably well conserved both across and within samples, and with no SNPs detected in the *pmoC* gene, indicating that this gene was highly conserved in the population. Only 10-37 base pair differences distinguish consensus sequences that were detected from each of the 13 samples. Using the reads that mapped to the genome from each sample, we called single nucleotide polymorphisms (SNPs) using the *inStrain* package [1]. As a result, between 14-160 SNPs were detected per sample, and those segregating variants were found on average in 70% of the samples, indicating that many polymorphisms were consistent between samples. We categorized each SNP as non-synonymous (NS) or synonymous based on predicted proteins. The overall ratio of non-synonymous SNPs to synonymous was 0.75. Across all samples, there are 23 genes with at least one non-synonymous SNPs (Supplementary Fig. 13a), 14 of them are hypothetical proteins with no domain detected. Within the nine with predicted function are two DNA polymerases and one endonuclease encoded by syntenic genes (i.e., genes _45, _46 and _47; Supplementary Fig. 13a).

To discern the population dynamics of individual variants over time, we tested for variants that significantly changed in frequency between 2016 and 2017 (z-test; $q < 0.05$). We found that variants that changed in frequency were about 2x as likely to be non-synonymous and that there were 14 non-synonymous variants that changed in frequencies between 2016 and 2017. These variants were found in 3 genes (Supplementary Fig. 13b), while two of these genes were unannotable, the third was an endonuclease with 3 amino acid variants that were present in most 2016 samples and absent from all but one 2017 sample. Overall, there was no average reduction or change in the average genetic diversity within the population between 2016 and 2017, indicating that any selective pressures present were not strong enough for selective sweeps of individual genotypes. Taken together, these results imply that the genes most quickly evolving in the phage population play largely unknown ecological roles.

**The confirmation of Methyloparacoccus_57 with low abundance in some BML samples**

When the biomass of a given population only accounts for a small fraction of that of a collected sample, *de novo* metagenomic assembly and subsequent analyses may not be able to detect the corresponding population with assembled fragments.

In this study, the host-phage relationship is predicted based on the similarity of the PmoC sequences (among those from pmoC-phages and bacterial methanotrophs) and followed by evaluation of the co-occurrence of phage and its predicted host (based on their genomic sequences assembled from metagenomic data). The pmoc-phages of BML_2, BML_3 and BML_3 were predicted with the potential host of Methyloparacoccus_57, however, assembled fragments of this population could only be detected in 14 of the 28 analyzed BML samples. To evaluate the existence of Methyloparacoccus_57 in the other 14 BML samples (from which the ribosomal protein S3 of Methyloparacoccus_57 was not assembled), we firstly curated the genome of Methyloparacoccus_57 (2,444,800 bp in length) from the sample of BML_10242017_9.75m (which has the highest sequencing coverage of this population), then the quality reads of all BML samples were individually mapped to the curated Methyloparacoccus_57 genome, with 2 mismatches allowed for each mapped read (> 98.6% nucleotide similarity). As expected, for the samples with Methyloparacoccus_57 fragments assembled, the number of reads (18,252 - 556,226 reads) mapped to a scaffold is strongly correlated to the length of the corresponding scaffold (sample names in black; Supplementary Fig. 6a). For the 14 BML samples without Methyloparacoccus_57 rpS3 assembled (sample names in red; Supplementary Fig. 6a), though only 380 - 6,046 reads were mapped to the curated Methyloparacoccus_57 genome, the number of reads mapped to a scaffold is also strongly correlated to the length of the corresponding scaffold. We also checked the reads mapped to the scaffold (i.e., BML_10242017_9_75m_scaffold_435) with the ribosomal proteins (Supplementary Fig. 6b), and found all samples with reads mapped to this region. *In summary, we concluded that Methyloparacoccus_57 was in all the 28 analyzed BML samples, though some of them with very low abundance.*

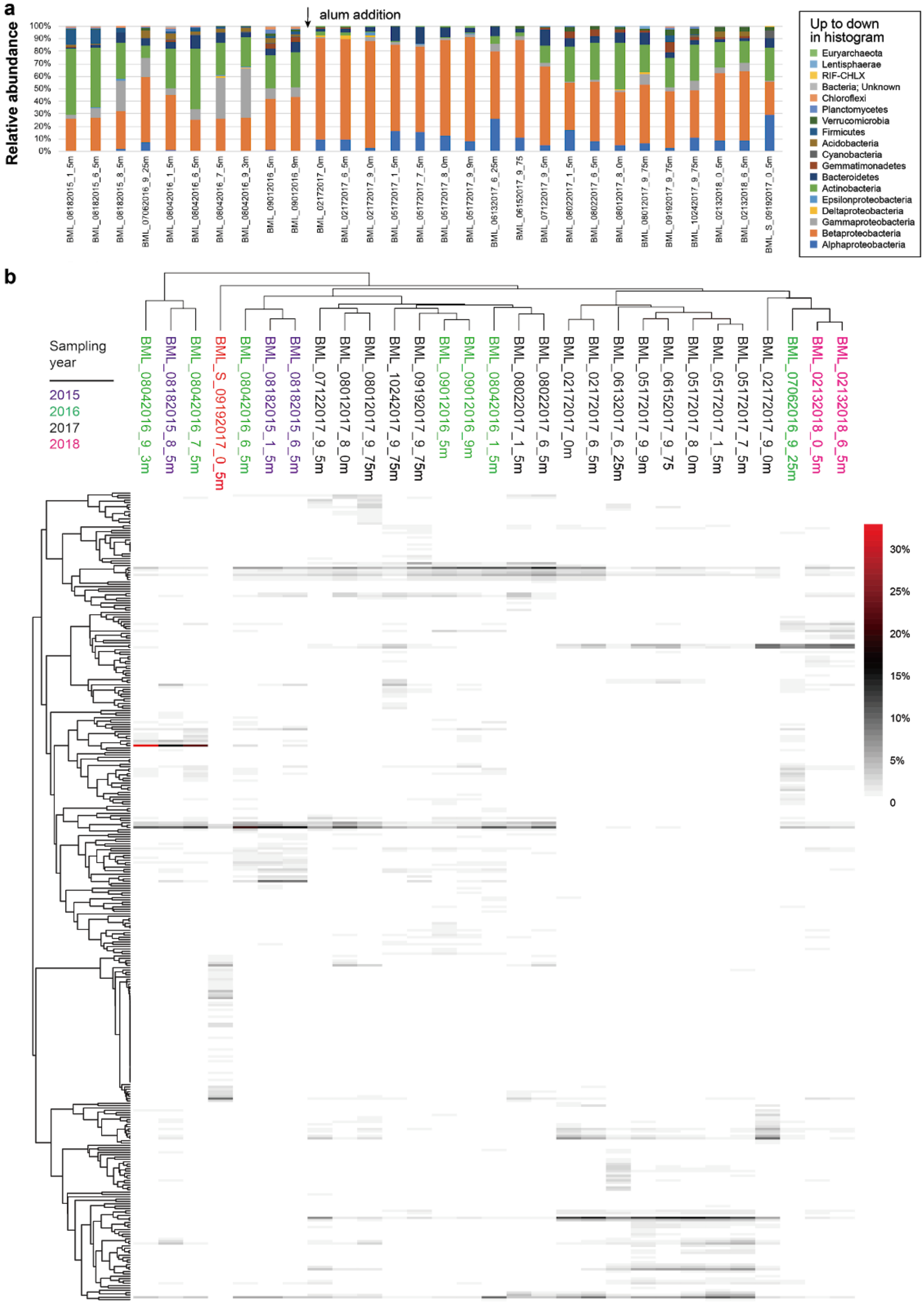


**Supplementary Fig. 1. Microbial community composition of BML and BML_S samples.** (a) The relative abundance of microbial phyla (or classes for Proteobacteria) in BML and BML_S samples. The analysis was performed based on ribosomal protein S3 (rpS3) genes (see methods in the main text for details). The addition of alum in 2016 was conducted to lower the available organic carbons in the water column. The communities were dominated by Actinobacteria, Alphaproteobacteria, and Betaproteobacteria. (b) The clustering analyses of BML and BML_S samples based on the relative abundance of microbial species/strain (determined based on rpS3) detected in the samples. Note that the BML_S microbial community is very different from the BML ones. The sampling year of samples is indicated by different colors. The clustering was performed using the R package of "pheatmap" [2] with the "correlation" clustering algorithm and "average" method.

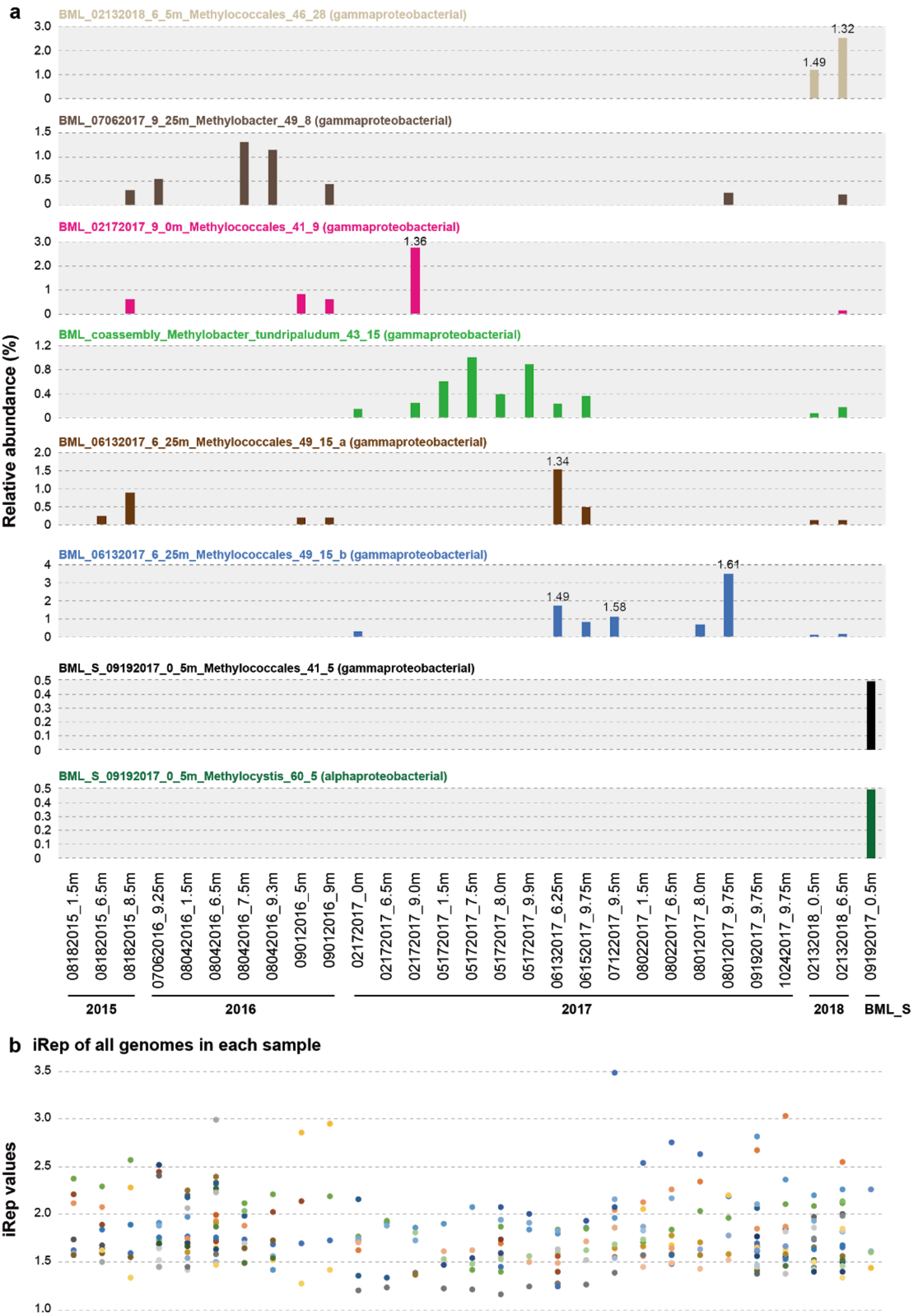


**Supplementary Fig. 2. The growth rate of microorganisms detected in BML and BML_S samples.** (a) The growth rate (iRep values) and relative abundance of alphaproteobacterial and gammaproteobacterial methanotrophs. The taxonomic information of methanotrophs are shown in the brackets, the iRep values are shown above the bars indicating relative abundance. (b) The growth rate (iRep values) of all microorganisms. Only those with at least a 5X coverage were calculated for growth rate via iRep analyses.

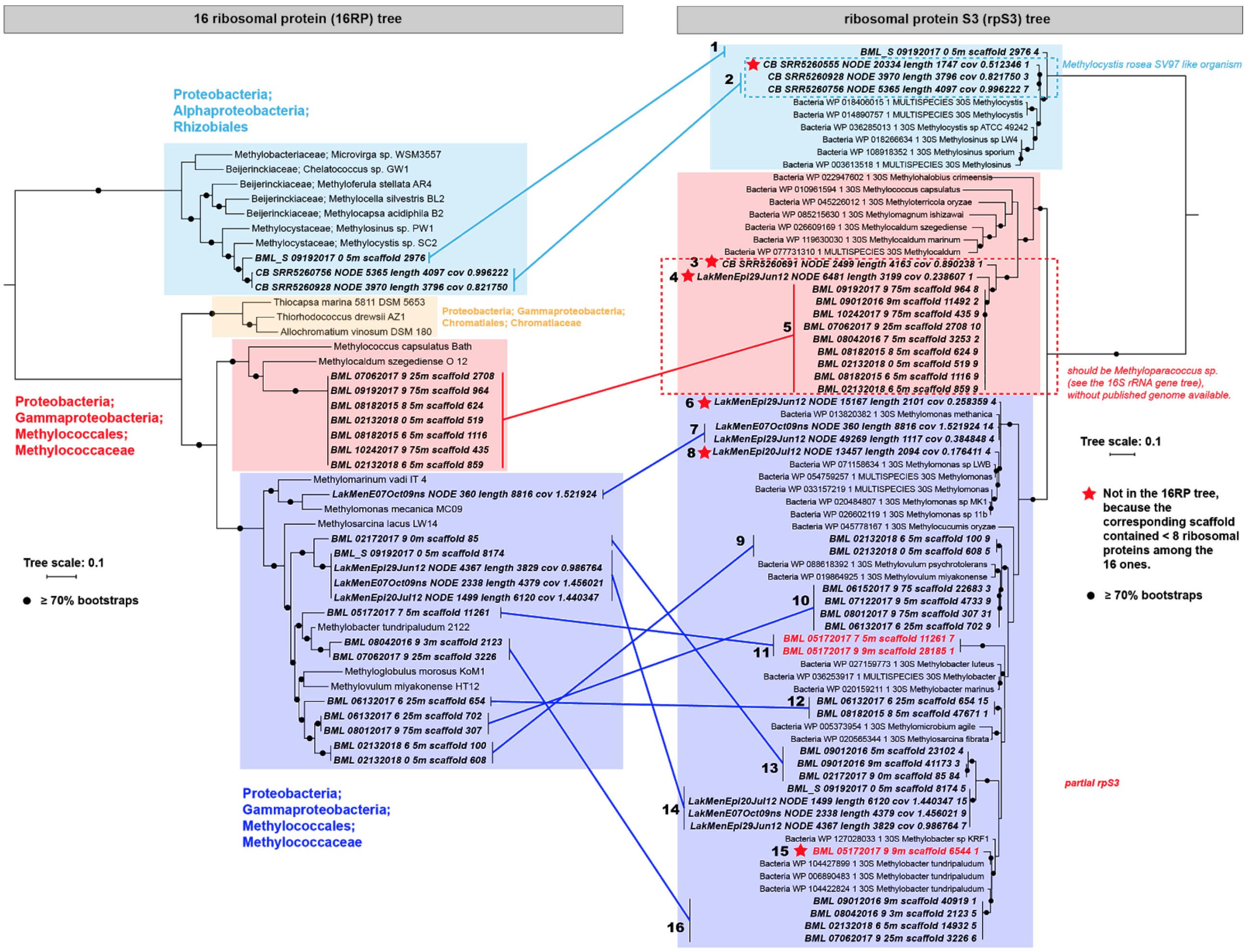


**Supplementary Fig. 3. Phylogenetic analyses of bacterial methanotrophs.** It was analyzed based on (left panel) 16 ribosomal proteins (16RPs), (right panel) ribosomal protein S3 (rpS3) and 16S rRNA (Supplementary Fig. 3 - continued). The scaffolds appeared in both trees are linked by lines and marked with a number for reference of the 16S rRNA tree (see next page for it). The scaffolds with 7 or fewer of the 16RPs were not included in the 16RPs and indicated by red stars in the rpS3 tree. The major lineages are highlighted in colored backgrounds. References were selected based on the rpS3 BLASTp search (top 5 hits for each). For a summary based on all three phylogenetic analyses: alphaproteobacterial methanotrophs were detected in CB and BML_S samples, gammaproteobacterial methanotrophs were detected in BML, BML_S, CB and LM samples. Note that the proteins and 16S rRNA genes were predicted from scaffolds with a minimum length of 1 kbp.

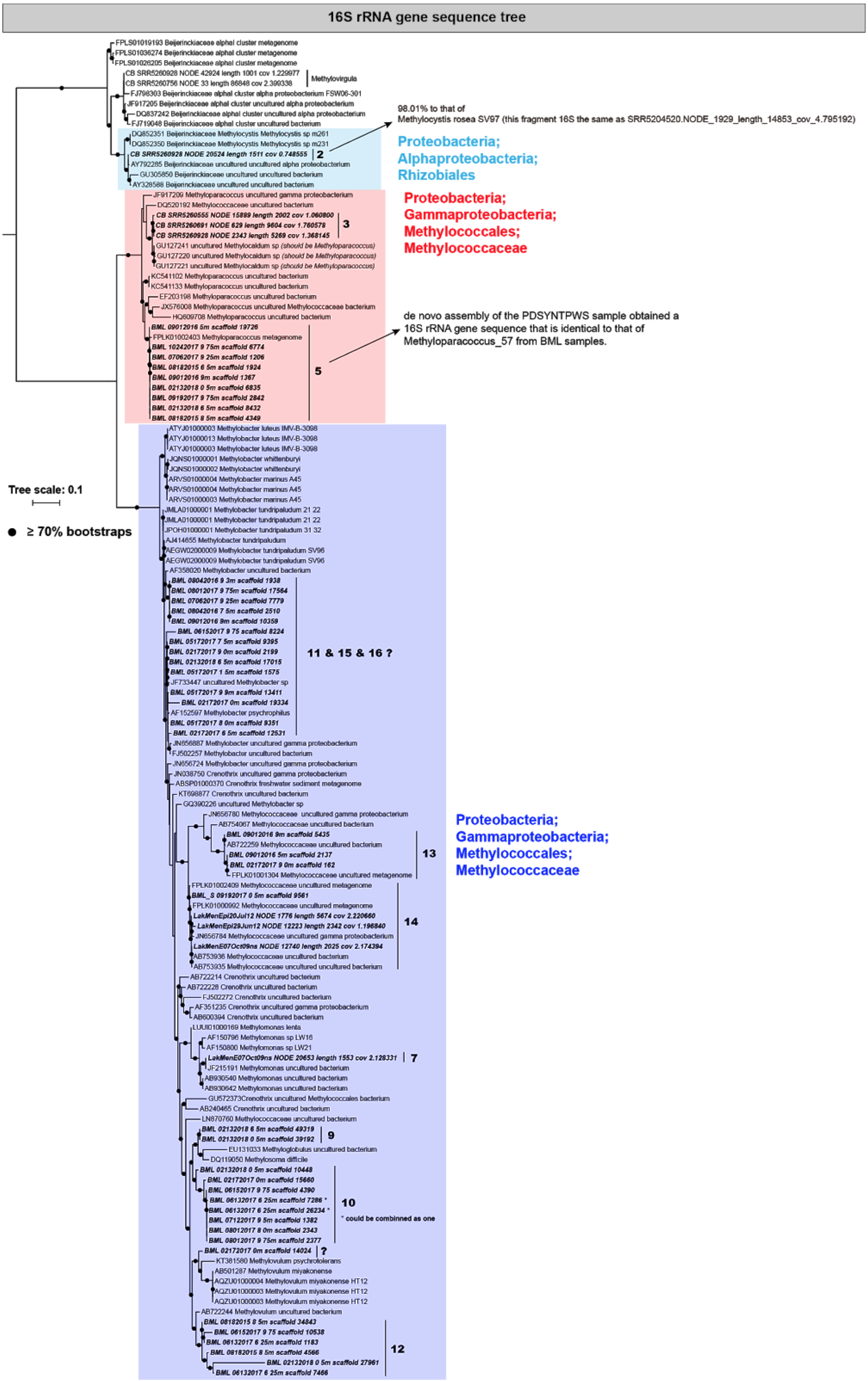


**Supplementary Fig. 3 - continued. Phylogenetic analyses of bacterial methanotrophs.** The 16S rRNA genes that could not be found with a match in 16RP or rpS3 tree are indicated a question mark.

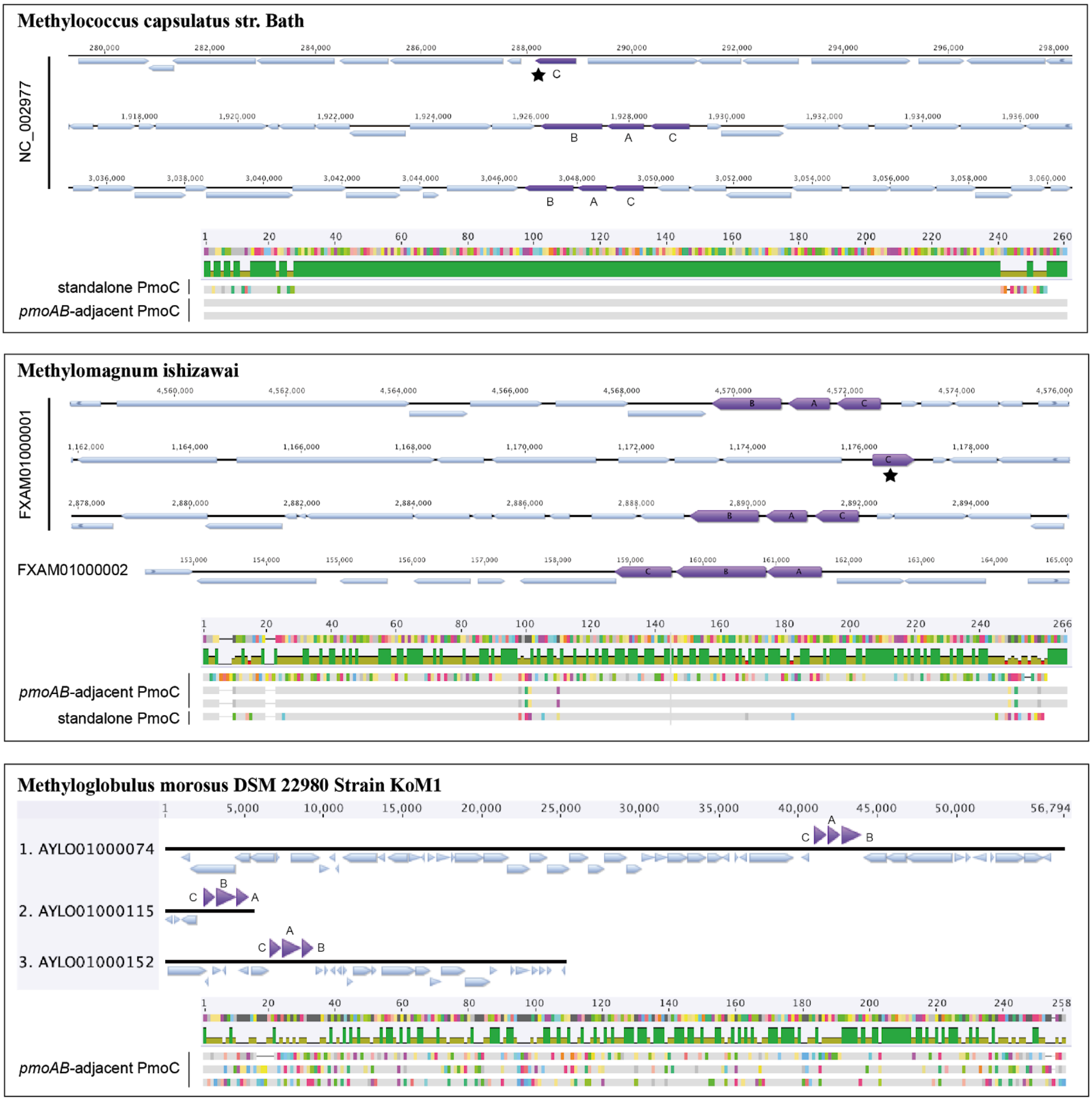


**Supplementary Fig. 4. The pMMO subunits detected in published bacterial methanotroph genomes.** The order of three subunits may vary from operon to operon (*pmoACB* or *pmoABC*). The protein sequence alignment of *pmoAB*-adjacent and standalone PmoC are shown at the bottom of each example. It is interesting that the PmoC from the same genome could be divergent from each other, and the standalone PmoC could be very similar to the ones within an operon. For analyses, the Genbank files of the genomes were downloaded from NCBI RefSeq, and the figures were drawn via Geneious with the pMMO subunits highlighted based on Genbank annotations. The standalone of *pmoC* genes are indicated by black stars.

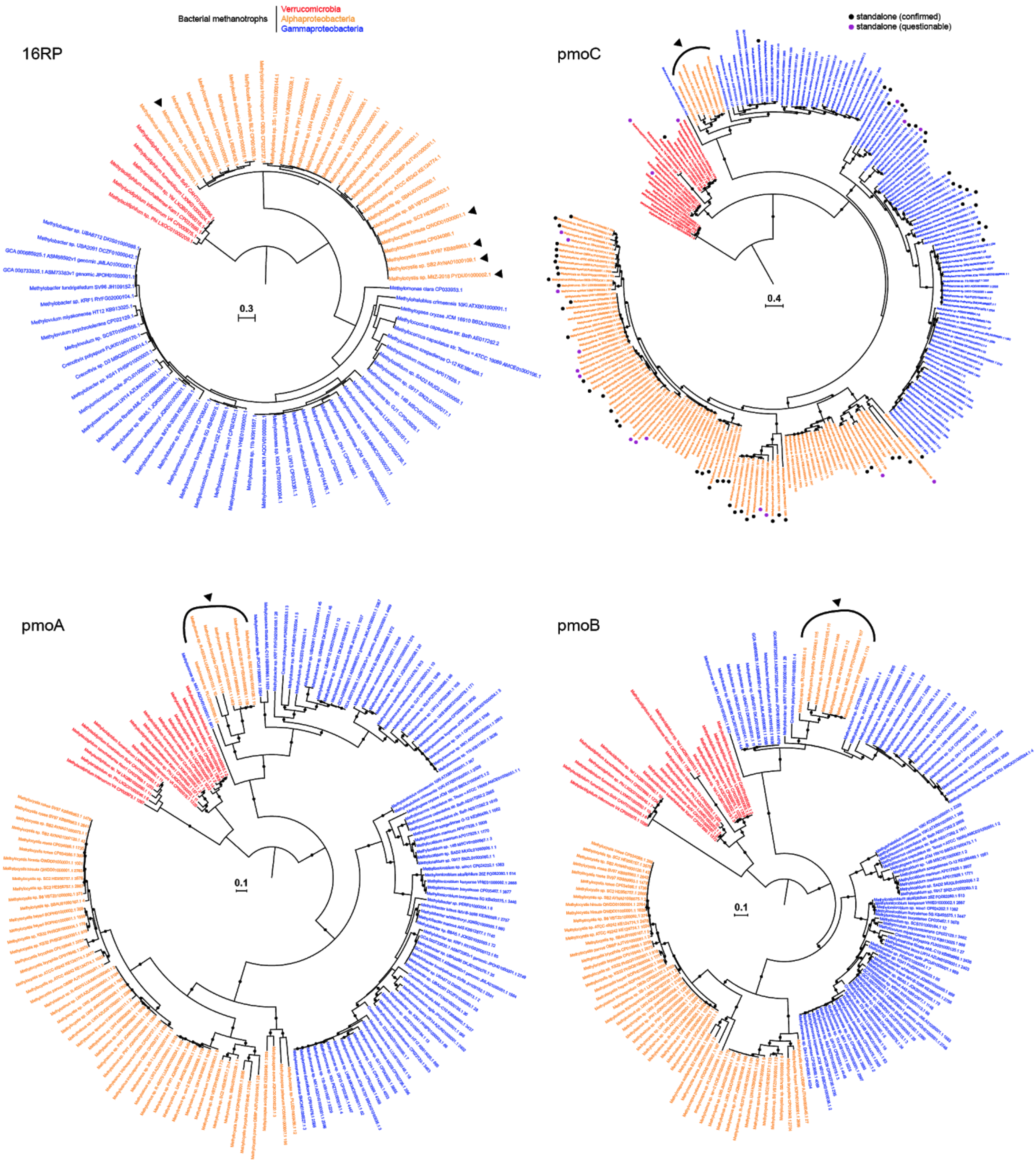


**Supplementary Fig. 5. Phylogenetic analyses of published bacterial methanotrophs based on concatenated sequences of 16 ribosomal proteins and pMMO subunits (PmoC, PmoA, and PmoB).** Only those genomes with a scaffold containing eight or more of the 16 ribosomal proteins are included in the 16RP-based phylogenetic analyses. The ones with one copy of their pMMOs phylogenetically related to some gammaproteobacterial methanotrophs are indicated by black triangles. Some bacterial methanotrophs genomes encode standalone *pmoC* (without *pmoA*/*pmoB* nearby), which are confirmed (via genetic context) and indicated by black solid circles, and those *pmoC* genes detected at the end of scaffolds are indicated by black empty circles.

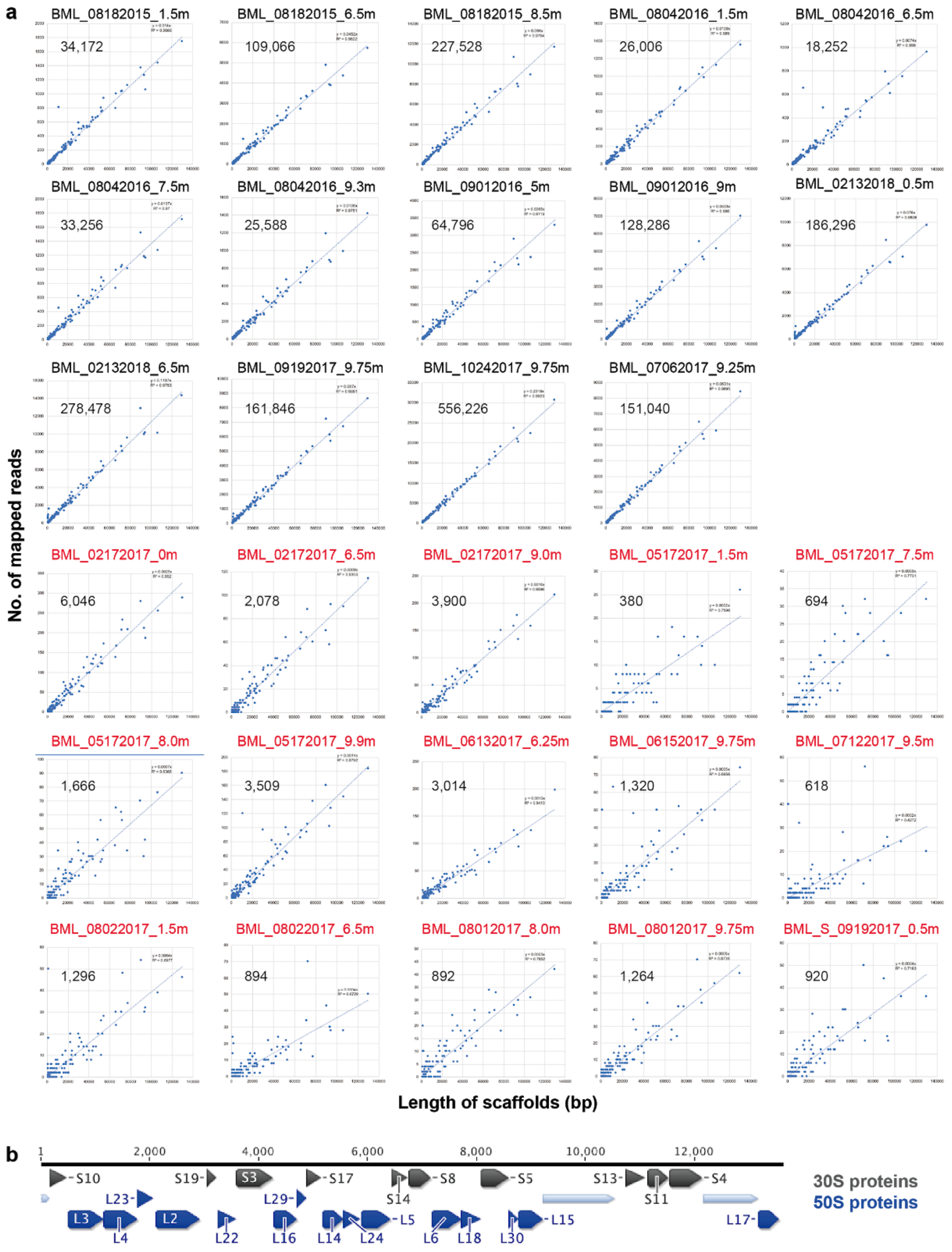


**Supplementary Fig. 6. The detection of Methyloparacoccus_57 with low abundance in some BML samples.** For each sample, the number of mapped reads to each scaffold was plotted as the function of the length of the corresponding scaffold. The total number of reads mapped to Methyloparacoccus_57 in each sample is shown. The sample names shown in black were detected with Methyloparacoccus_57 via assembled scaffolds and reads mapping, the ones are shown in red via only reads mapping. See Supplementary information (section "The confirmation of Methyloparacoccus_57 with low abundance in some BML samples") for details.

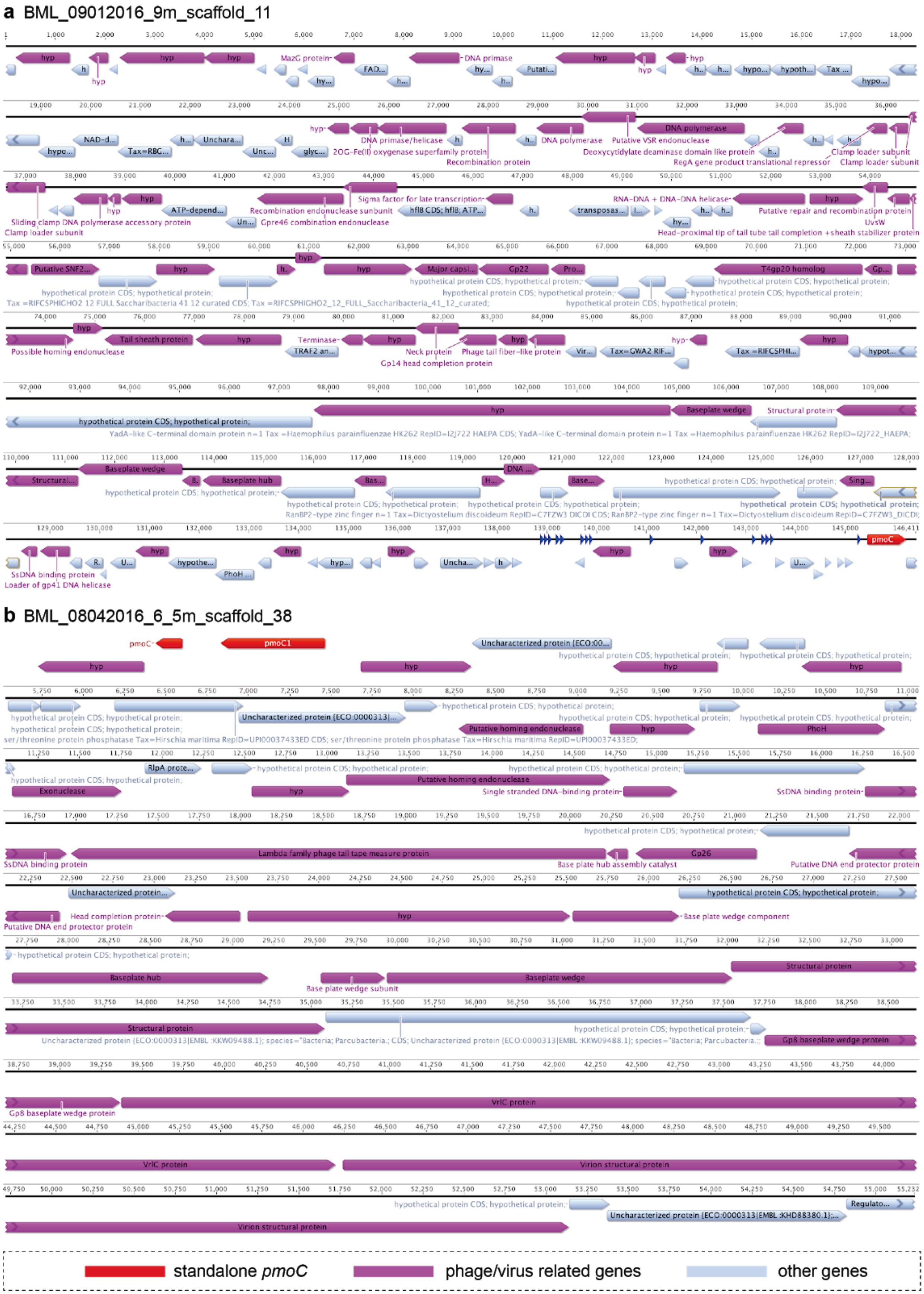


**Supplementary Fig. 7. Examples of *pmoC* gene detected on scaffolds with phage signals.** The phage/virus related genes were identified by the search against the KEGG, UniRef and UniProt databases (see methods in the main text for details).

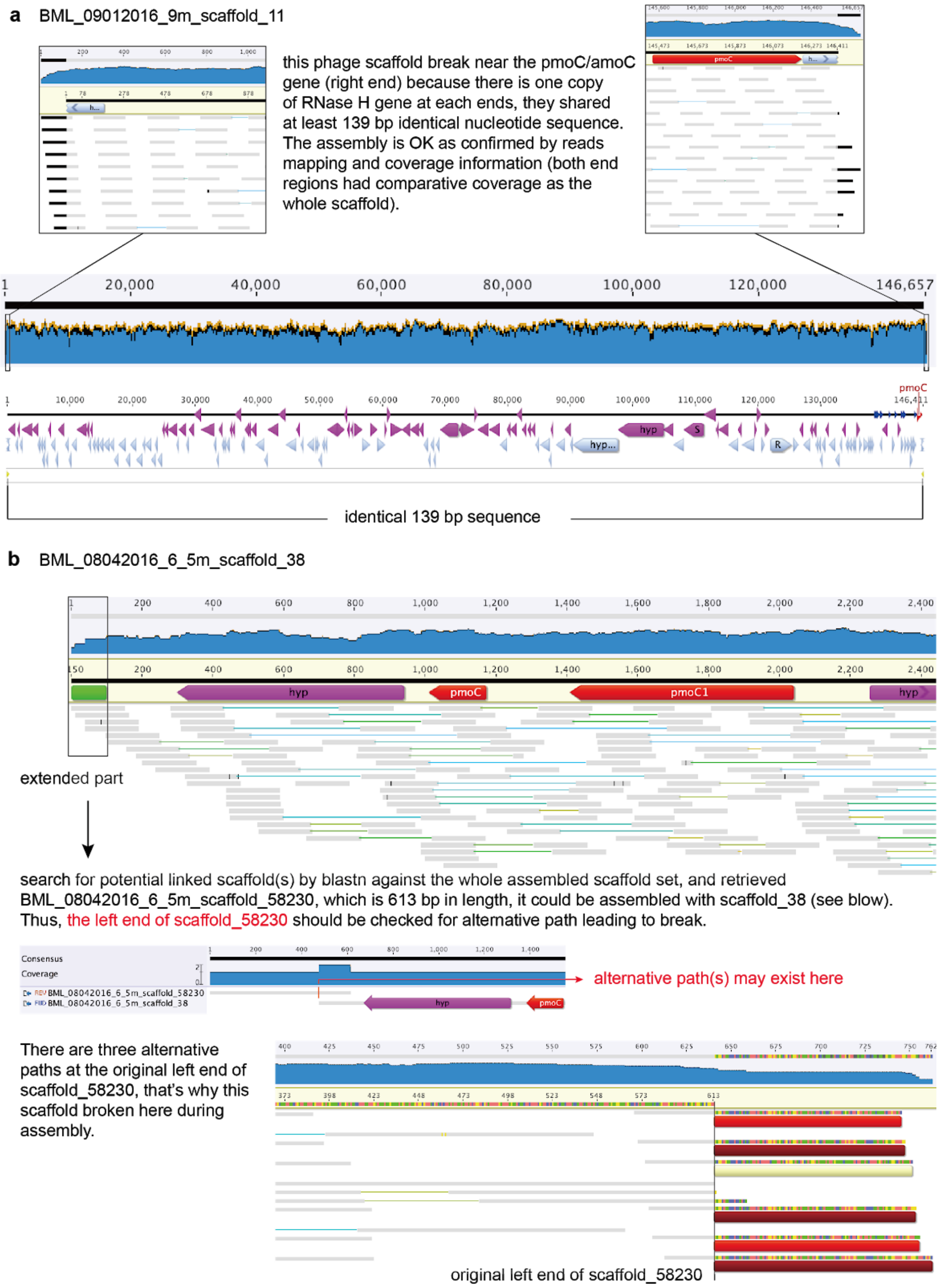


**Supplementary Fig. 8. Examples showing the reasons why phage-related scaffolds broke at or near the *pmoC* gene.** In detail, the example (a) is due to the repeat sequences at the two ends of the scaffolds, (b) is due to the existence of alternative paths for the part that scaffold_38 could be joined with.

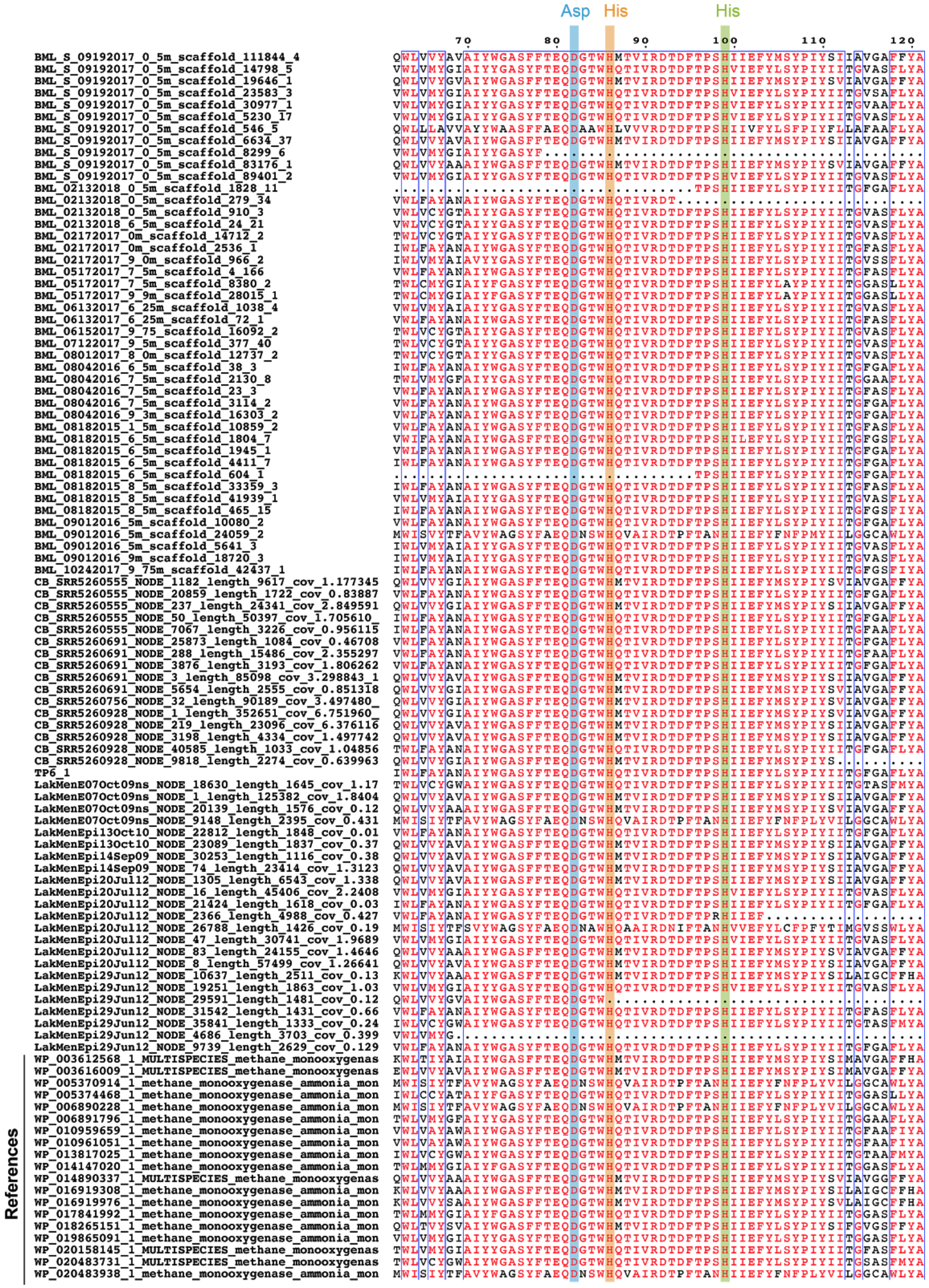


**Supplementary Fig. 9. Alignment of bacterial and phage-associated PmoC sequences detected in BML, BML_S, LM and CB samples.** The residues involved in $O_2$ binding and methane oxidation are highlighted.

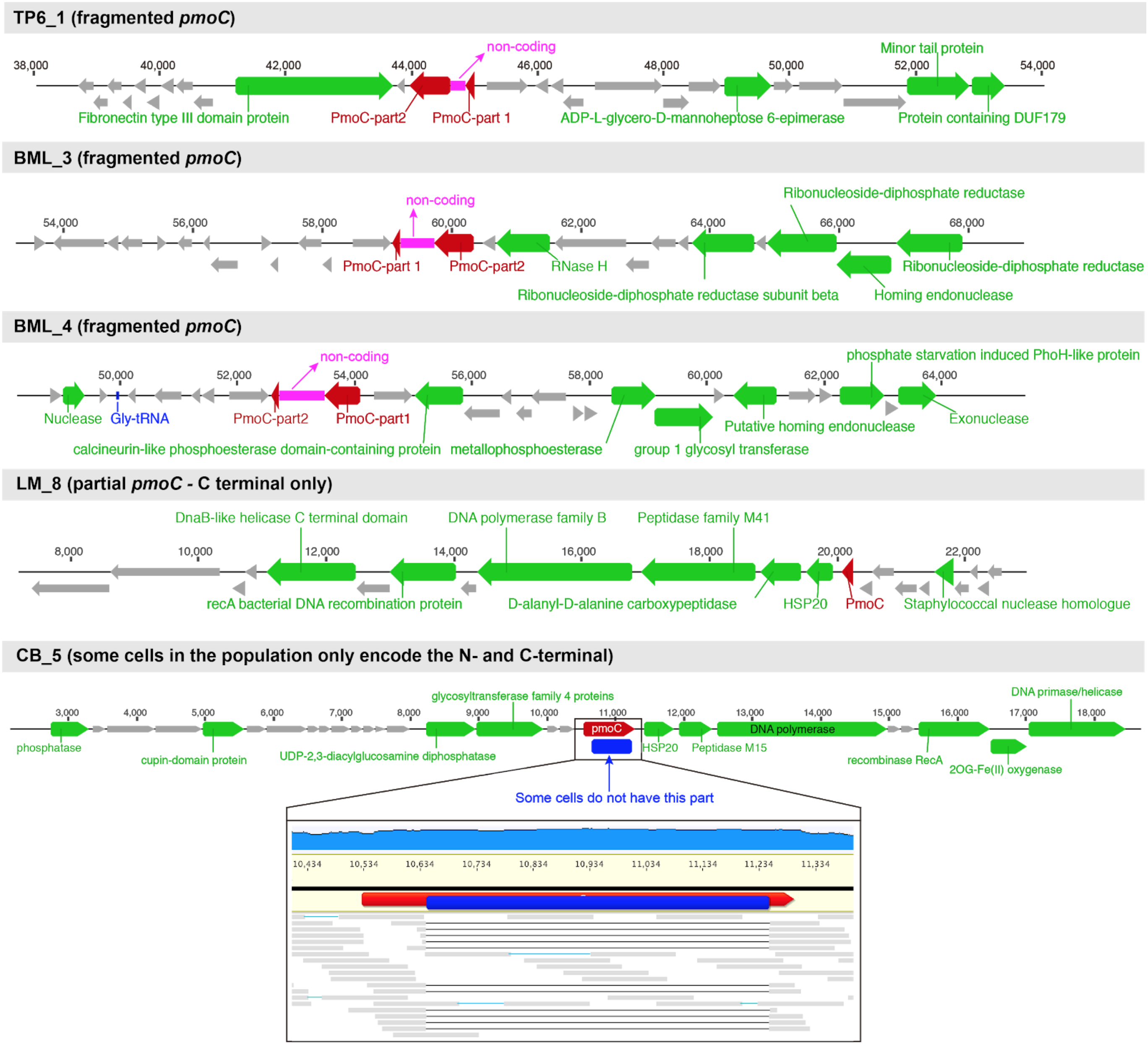


**Supplementary Fig. 10. Genetic context of fragmented and partial *pmoC* genes in pmoC-phages.** Fragmented or partial phage-associated PmoC were detected in pmoC-phages with predicted hosts of alphaproteobacterial methanotroph (i.e., LM_8, CB_5) and gammaproteobacterial methanotrophs (i.e., TP6_1, BML_3 and BML_4). The protein-coding genes with functional annotation are shown in green, the ones without functional annotation in grey, tRNA in blue. Some CB_5 cells contained only part of the *pmoC* gene, which is shown in detail.

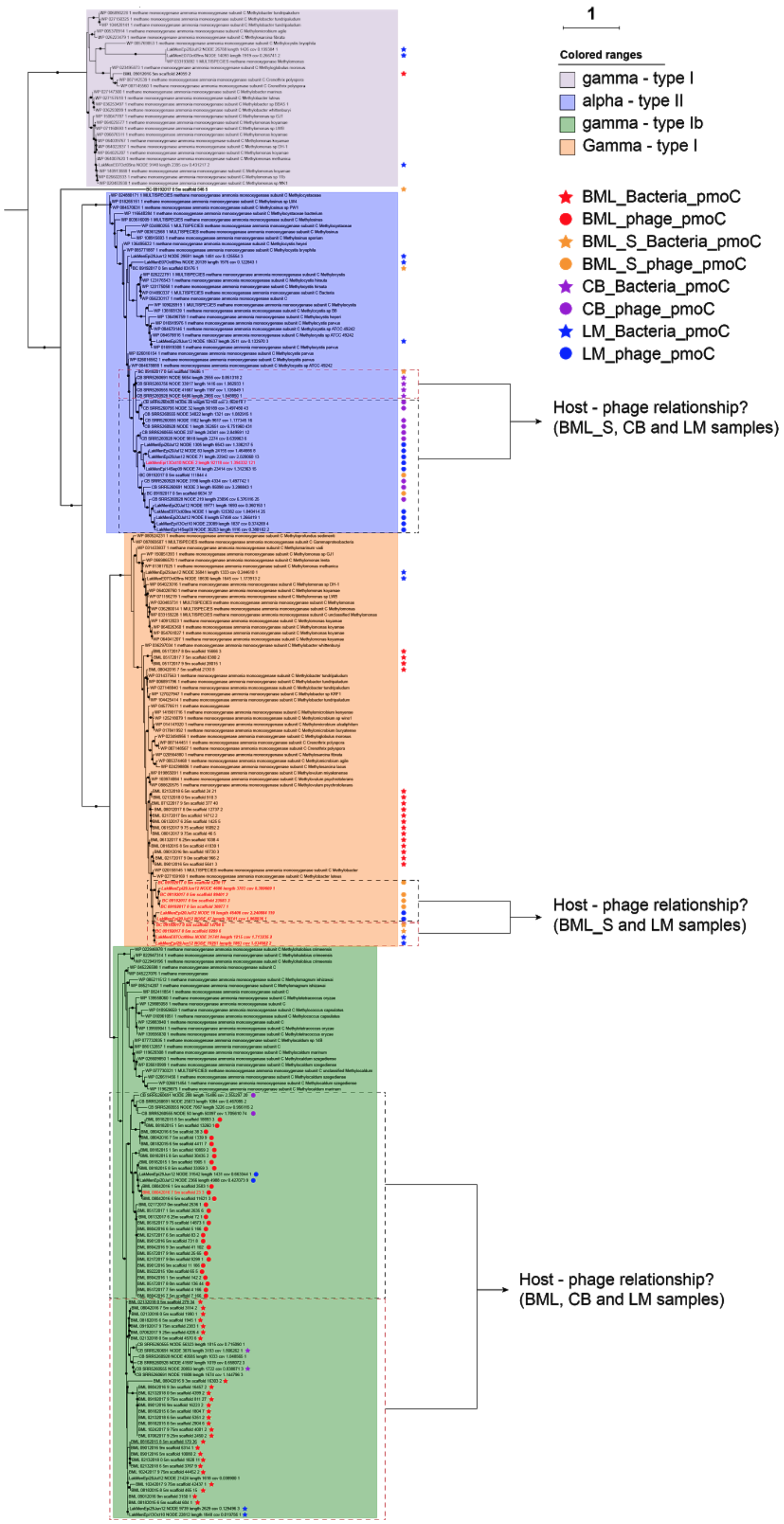


**Supplementary Fig. 11. The full phylogenetic tree of bacterial and phage-associated PmoC detected in BML, BML_S, LM, CB and TBL samples.** PmoC from published bacterial methanotrophs with genomes available are included for reference, see Supplementary Tables 3 and 6 for details. Please note that only some metagenomic datasets from LM and CB have been re-analyzed and PmoC were included here.

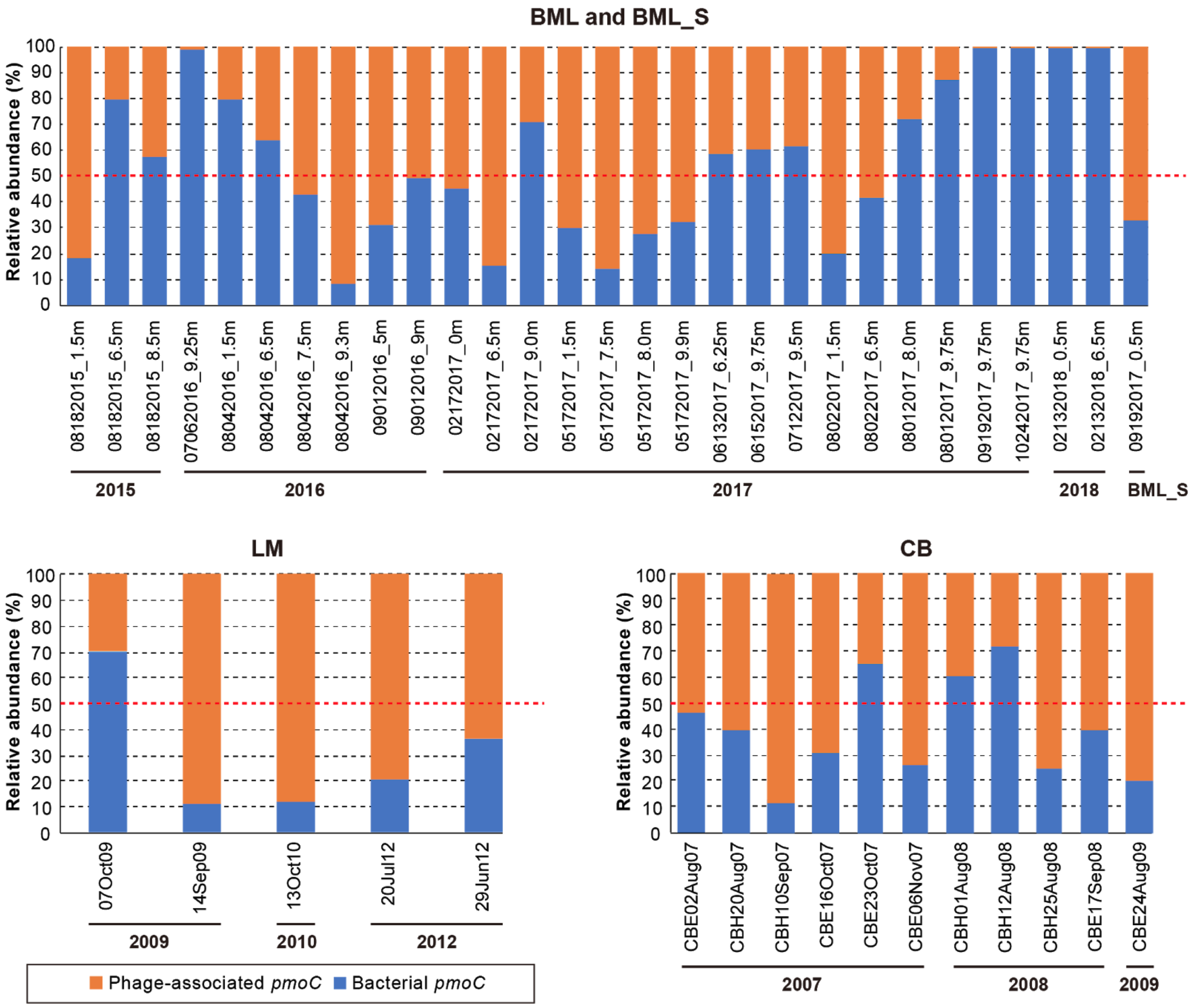


**Supplementary Fig. 12.** The cumulative relative abundance of bacterial and phage-associated *pmoC* genes in samples from BML, BML_S, LM, and CB. Please note that only some of the metagenomic datasets from LM and CB have been re-analyzed, for which ones the data are shown here. The quality reads from each sample were mapped to the representative scaffolds with the detected bacterial or phage-associated *pmoC* gene, and the sequencing coverage was calculated for each scaffold, the cumulative coverage of all bacterial pmoC scaffolds was summed as *i*, and that of phage-associated pmoC scaffolds summed as *j*, the cumulative relative abundance of bacterial *pmoC* was calculated as $i/(i + j)$ X 100%, and that of phage-associated *pmoC* was calculated as $j/(i + j)$ X 100% (or 1 - $i/(i + j)$ X 100%).

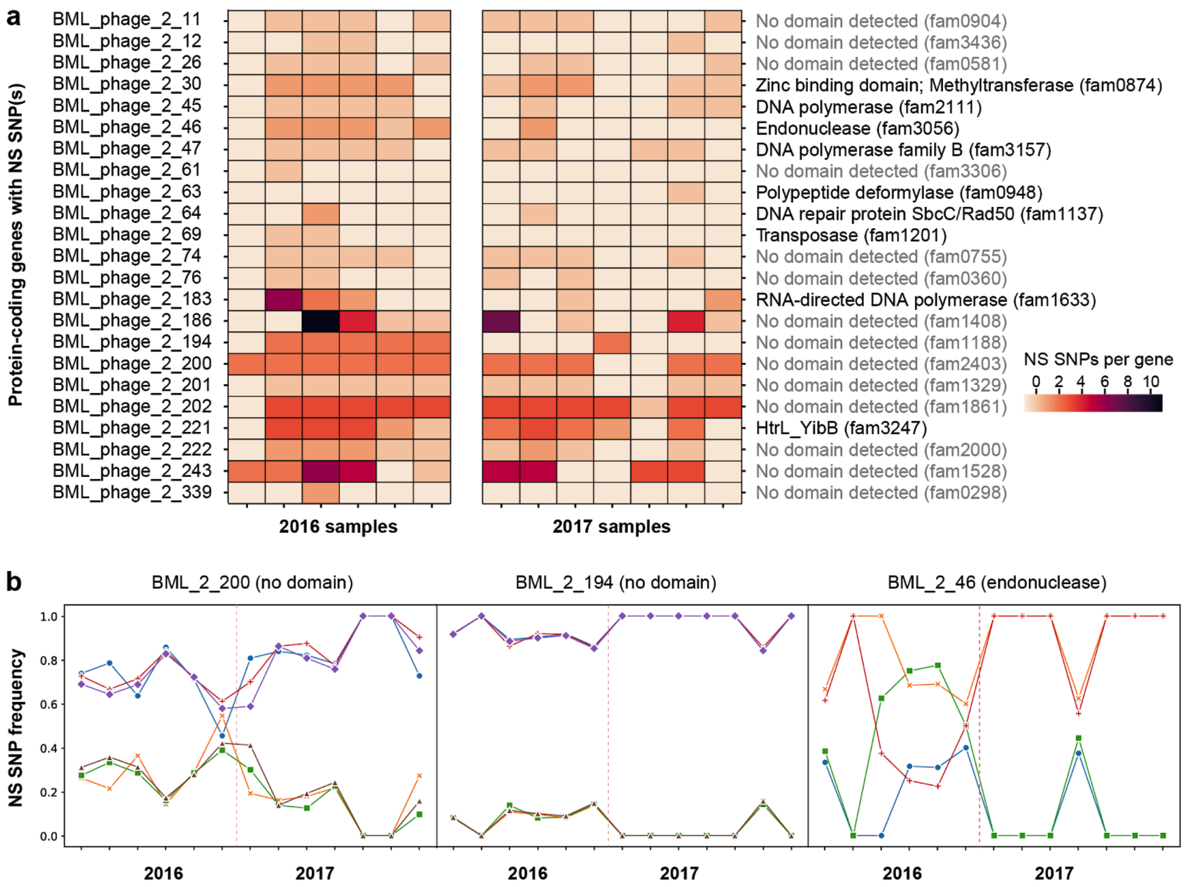


**Supplementary Fig. 13. The SNPs analyses of pmoC-phage BML_2. (a) The number of SNPs detected in protein-coding genes and their functional annotation.** Only protein-coding genes with non-synonymous (NS) SNPs are shown and listed in the order of genes on the chromosome. The annotation of protein-coding genes and their corresponding protein families are listed on the right. (b) Frequencies of NS SNPs that changed significantly (z-test; q<0.05) between 2016 and 2017 samples. Each allele and its corresponding alternative allele are represented by a line, and SNPs are grouped by the origin of the genes.

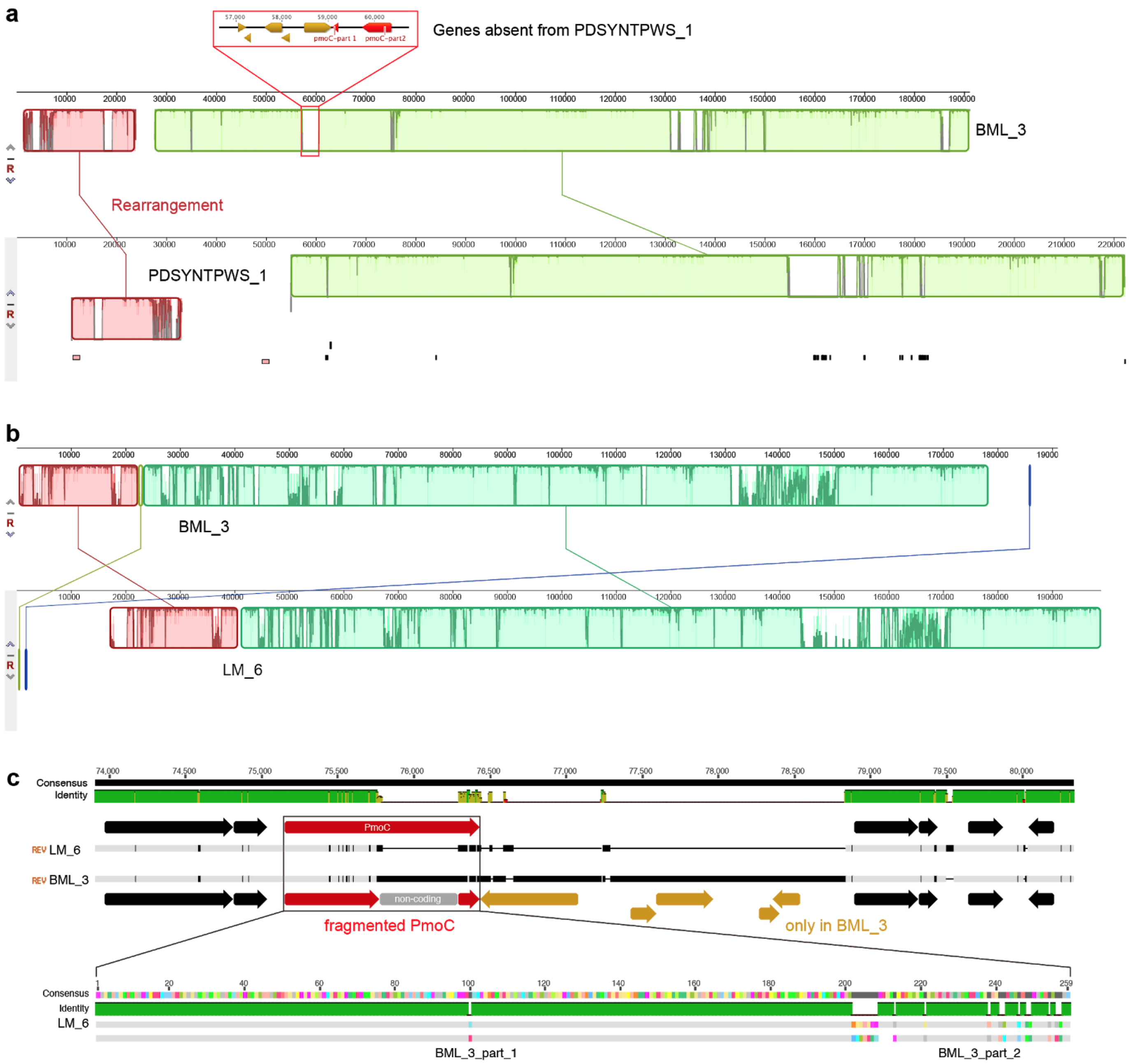


**Supplementary Fig. 14 | Whole-genome alignment of BML_3, PDSYNTPWS_1 and LM_6.** (a)Mauve genome alignment viewer of BML_3 and PDSYNTPWS_1. The pmoC gene and five syntenic genes (all for hypothetical proteins) that present in BML_3 while absent from PDSYNTPWS_1 are shown in detail at the top. (b) Mauve genome alignment viewer of BML_3 and LM_6. (c) Genome alignment of the pmoC regions of LM_6 and BML_3. Five protein-coding genes near the fragmented PmoC are only present in BML_3. The alignment of PmoC sequences shows the breakpoint of the PmoC in BML_3.

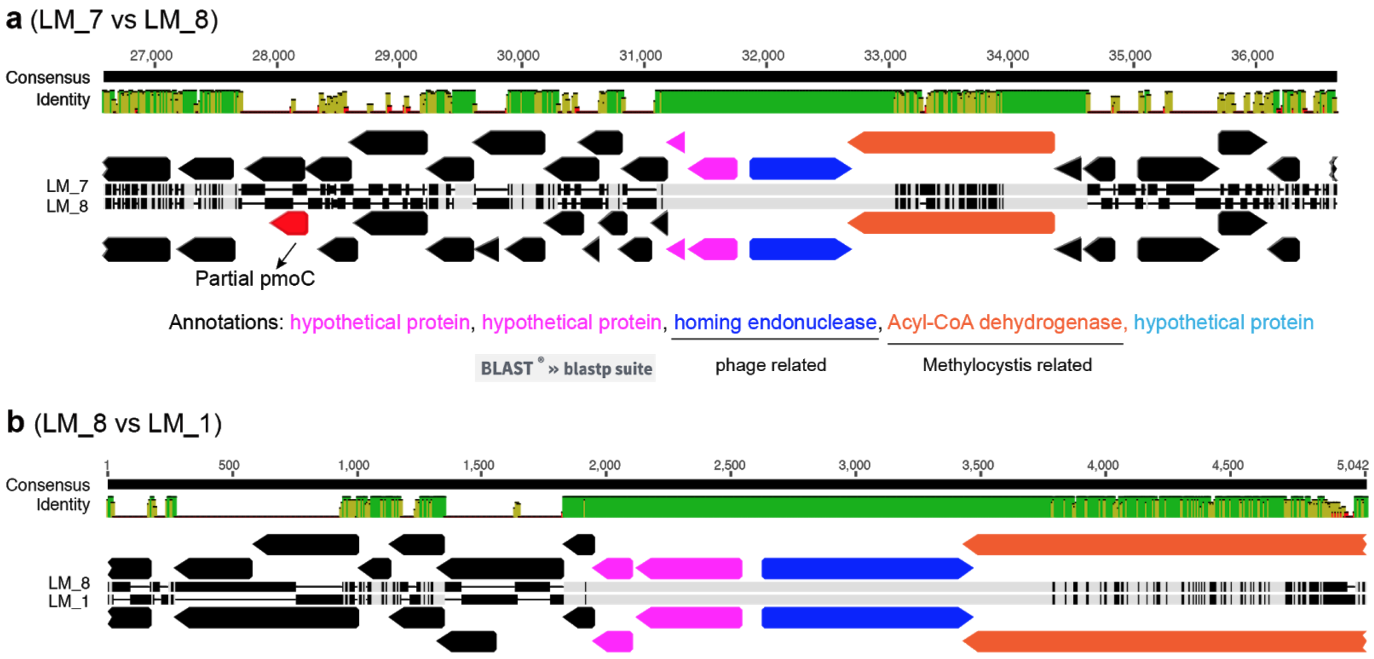


**Supplementary Fig. 15 | The shared region among LM_1, LM_7 and LM_8.** Manual curation and check were performed to confirm that these phage genomes share this region. The NCBI BLASTp information is shown for the two protein-coding genes that could be annotated.

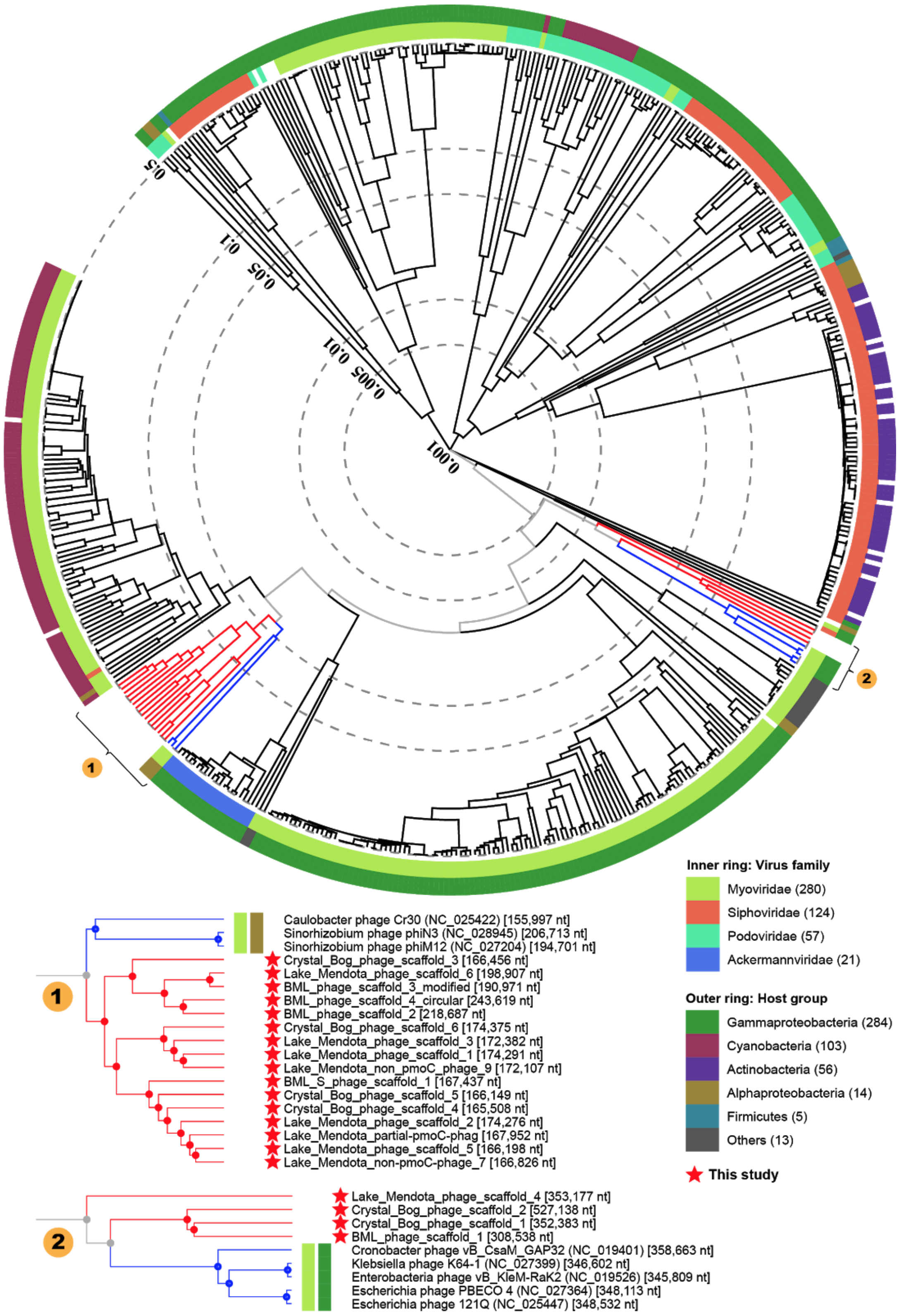


**Supplementary Fig. 16. The selection of reference phage/virus genomes based on Viptree analyses.** The analysis was performed by uploading the curated phage genomes reconstructed in this study via the viptree online analysis tool (https://www.genome.jp/viptree/), which generated the circular proteomic tree. Based on this, the reference virus/phage genomes with similar protein profiles with phages reported in this study are shown in detail at the bottom of the Figure. These reference genomes are used for protein families and phylogenetic analyses that are detailed in the main text.

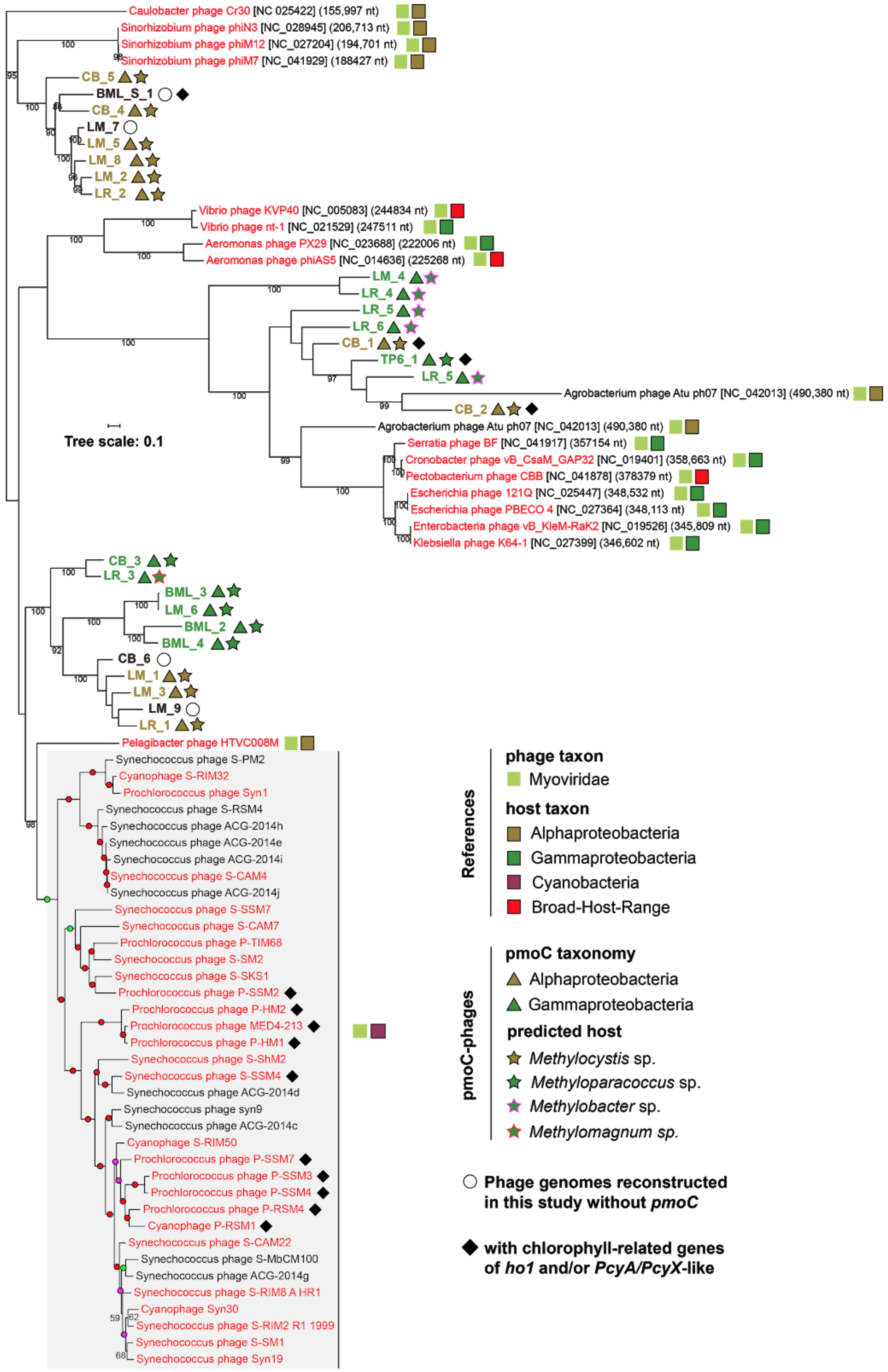


**Supplementary Fig. 17. Phylogenetic analyses of phages with genomes reported in this study based on DNA polymerases.** The taxa of reference phages and their hosts are shown with information from the NCBI virus-host database [3].

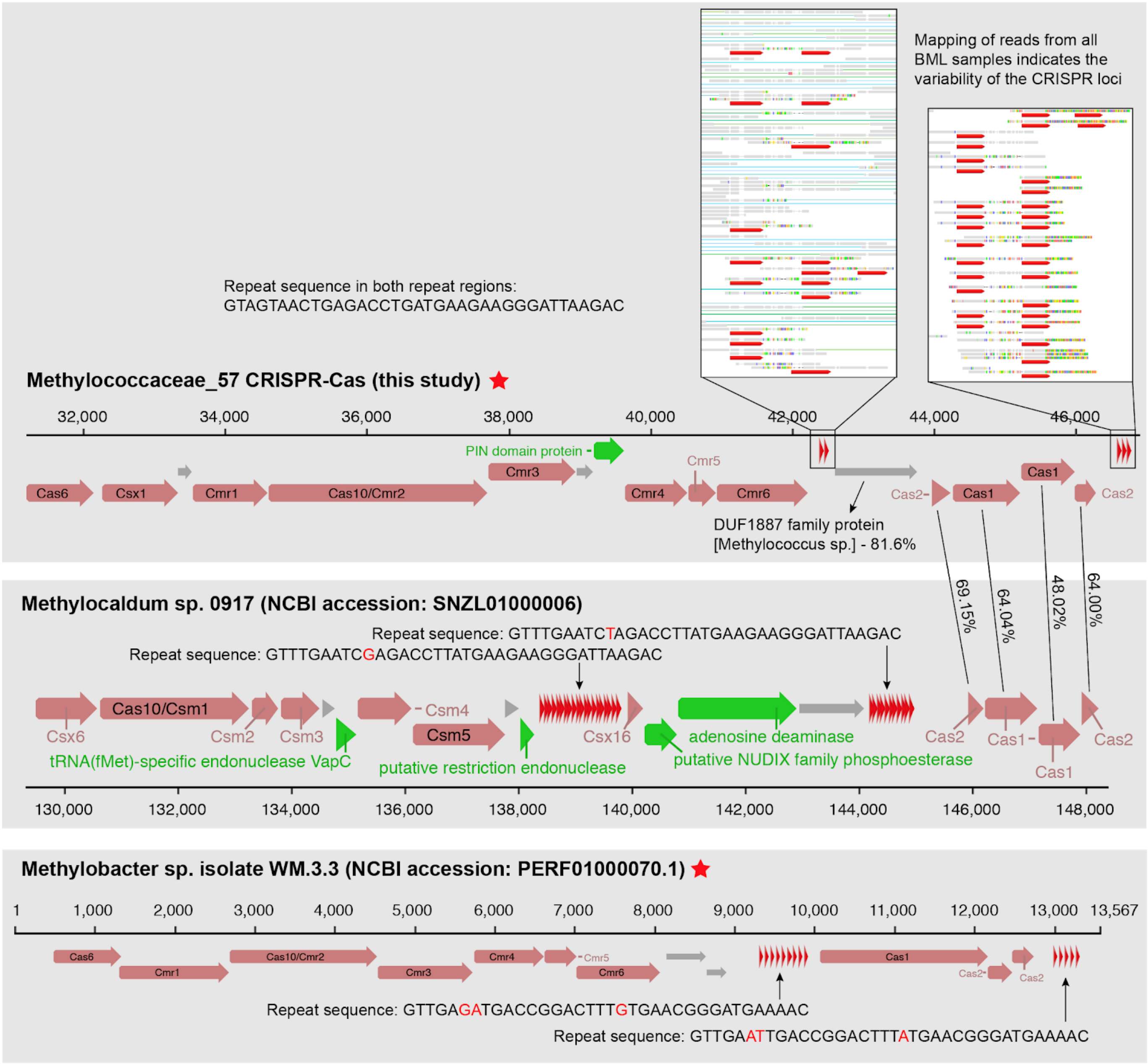


**Supplementary Fig. 18. CRISPR-Cas analyses of bacterial methanotrophs reported in this study and that already published.** The CRISPR-Cas system of Methyloparacoccus_57 (top panel) contains two repeat regions with the same repeat sequences, the Cas1 and Cas2 protein sequences are most similar to those from Methylocaldum sp. 0917 (middle panel), which also have two repeat regions and the repeat sequences are only one base divergent. The mapping of reads from all BML samples to the CRISPR scaffolds indicates divergences in spacer sequences, and we found one of the spacers matches the genomic sequence of pmoC-phage BML_4. Interestingly, one spacer from the published Methylobacteria sp. isolate WN.3.3 (bottom panel) also targets pmoC-phage BML_4. The CRISPR-Cas system of Methylobacteria sp. isolate WN.3.3 is also similar to that of Methyloparacoccus_57 by sharing the same type of other cas proteins excluding cas1 and cas2.

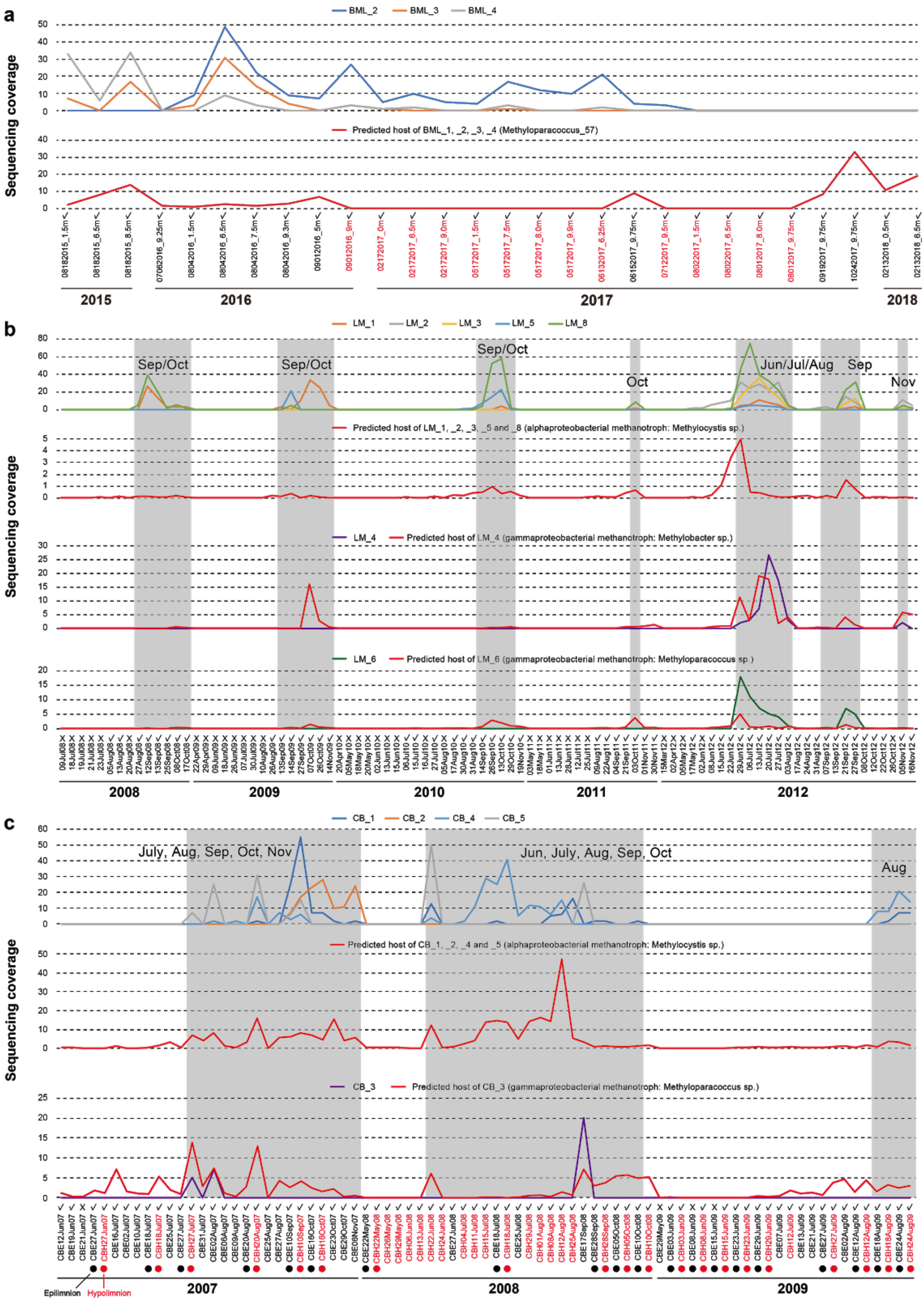


See next page for legend

**Supplementary Fig. 19. The sequencing coverage profiles of genomes of pmoC-phages and their predicted hosts in (a) BML, (b) Lake Mendota and (c) Crystal Bog samples.** The published pmoC-phage of BML_1 sampled in August 2011 from 0-10 cm of an oil sands lake [4] is predicted to have the same host as other BML pmoC-phages. The names of samples detected with very low abundances of Methyloparacoccus_57 are shown in red (see Supplementary Fig. 6 and Supplementary Information for details). Grey shading indicates times when both pmoC-phages and the predicted host are relatively abundant. When one bacteria was predicted as the host of multiple pmoC-phages, their profiles are shown in separate panels. The epilimnion and hypolimnion samples collected on the same day from Crystal Bog are paired and indicated by solid black and red circles, respectively. Only epilimnion samples were collected from Lake Mendota. A "✓" indicated the detection of bacterial *pmoC* gene(s) in the sample, while "X" indicates no detection.

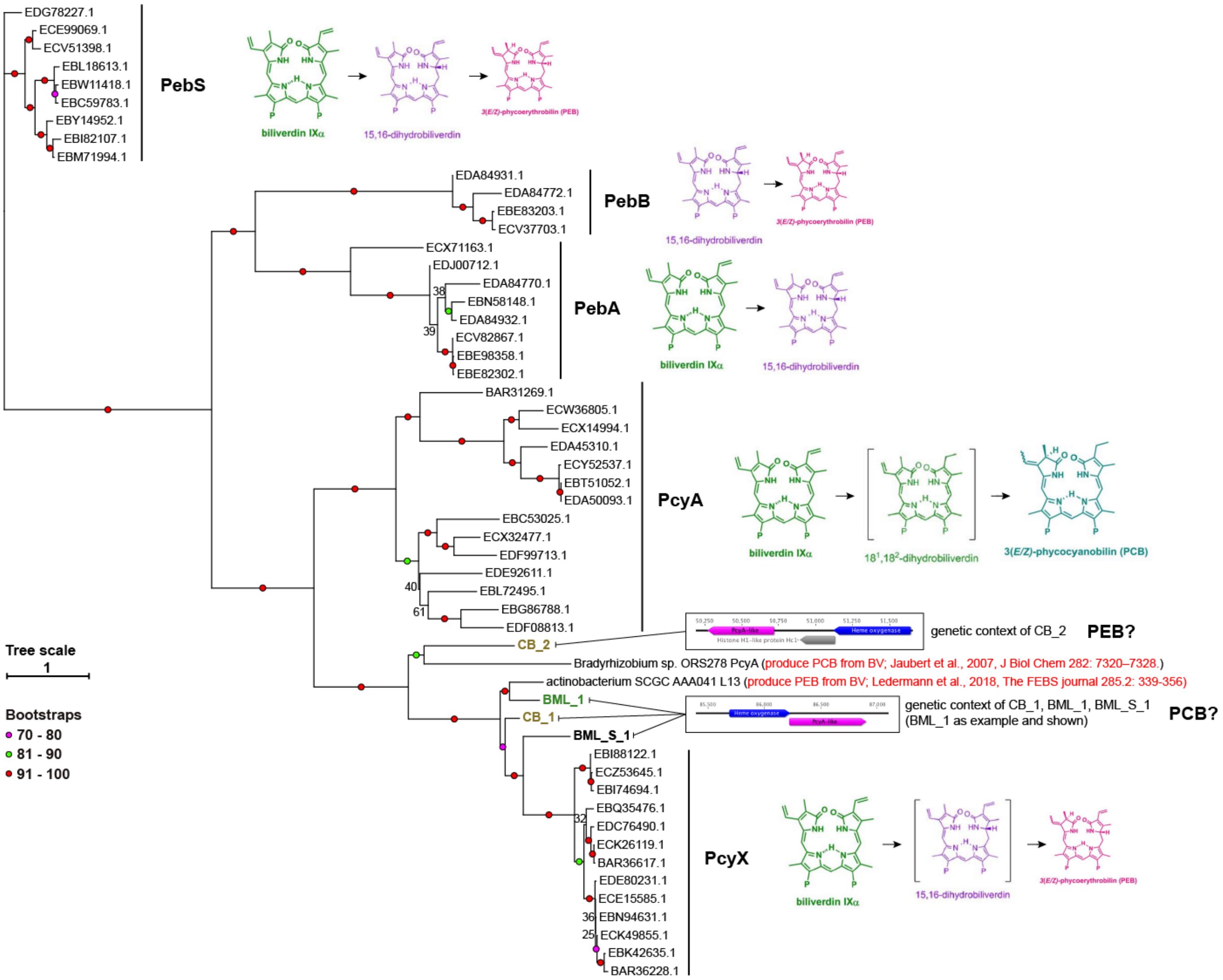


**Supplementary Fig. 20. Phylogenetic analyses of the Pcy-A like proteins encoded by three pmoC-phages and one non-pmoC-phage reported in this study.** The ferredoxin-dependent bilin reductases (FDBRs) reference sequences were retrieved from a previous study [5], along with the information of products from the corresponding proteins. The products of PcyA_Brady and PcyX_actino have been documented and reported [6], based on which the products by similar phage proteins reported in this study were speculated and shown. The genetic context of genes in phages reported in this study is shown in detail. The analyses were performed by firstly aligning the proteins using Muscle [7] and filtering the alignment by trimAl [8] to remove those columns with ≥ 90% gaps, followed by tree building with IQtree [9] using the “LG+G4” model.

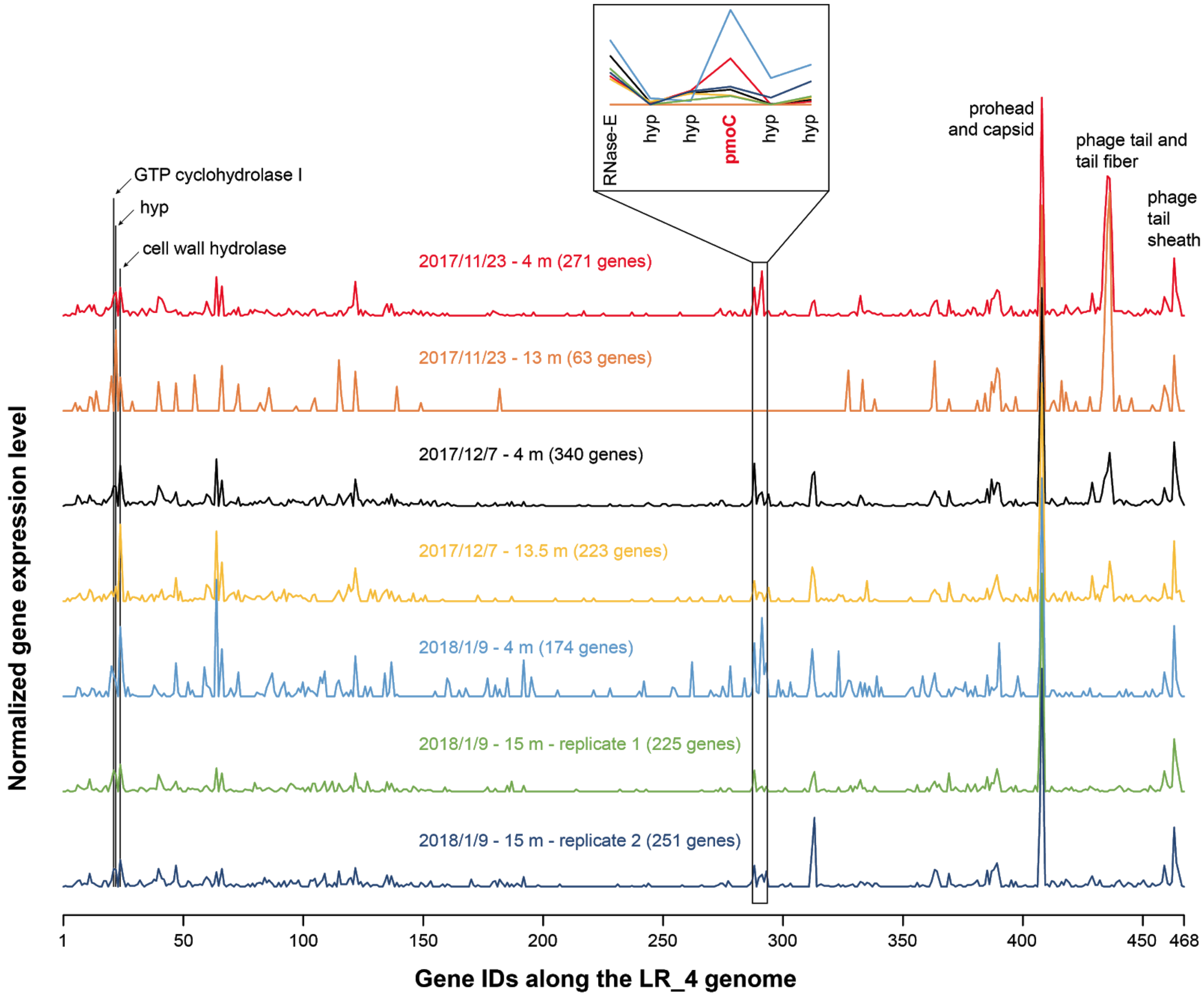


**Supplementary Fig. 21. The gene expression of pmoC-phages with genomes reconstructed from Lake Rotsee, with LR_4 as an example.** The sample collecting time points are shown for each sample above the gene transcriptional profile, followed by the sampling depth, and the number of genes transcribed in the brackets. Some genes with high transcriptional activities are highlighted with their annotations, the transcriptional levels of six syntenic genes including the *pmoC* genes are zoomed-in in the inserted figure.
